# Atypical response inhibition and error processing in 22q11.2 Deletion Syndrome and Schizophrenia: Towards neuromarkers of disease progression and risk

**DOI:** 10.1101/2020.04.29.068932

**Authors:** Ana A. Francisco, Douwe J. Horsthuis, Maryann Popiel, John J. Foxe, Sophie Molholm

## Abstract

22q11.2 deletion syndrome (also known as DiGeorge syndrome or velo-cardio-facial syndrome) is characterized by increased vulnerability to neuropsychiatric symptoms, with approximately 30% of individuals with the deletion going on to develop schizophrenia. Clinically, deficits in executive function have been noted in this population, but the underlying neural processes are not well understood. Using a Go/No-Go response inhibition task in conjunction with high-density electrophysiological recordings (EEG), we sought to investigate the behavioral and neural dynamics of inhibition of a prepotent response (a critical component of executive function) in individuals with 22q11.2DS with and without psychotic symptoms, when compared to individuals with idiopathic schizophrenia and age-matched neurotypical controls. Twenty-eight participants diagnosed with 22q11.2DS (14-35 years old; 14 with at least one psychotic symptom), 15 individuals diagnosed with schizophrenia (18-63 years old) and two neurotypical control groups (one age-matched to the 22q11.2DS sample, the other age-matched to the schizophrenia sample) participated in this study. Analyses focused on the N2 and P3 no-go responses and error-related negativity (Ne) and positivity (Pe). Atypical inhibitory processing was shown behaviorally and by significantly reduced P3, Ne, and Pe responses in 22q11.2DS and schizophrenia. Interestingly, whereas P3 was only reduced in the presence of psychotic symptoms, Ne and Pe were equally reduced in schizophrenia and 22q11.2DS, regardless of the presence of symptoms. We argue that while P3 may be a marker of disease severity, Ne and Pe might be candidate markers of risk.

## INTRODUCTION

22q11.2 Deletion Syndrome (22q11.2DS), otherwise known as DiGeorge or velo-cardio-facial syndrome (VCFS), is often characterized by relatively severe physical, cognitive and psychiatric manifestations (Shprintzen, 2008). Among the latter is a substantially increased risk for psychosis: A deletion on the long arm of chromosome 22 confers one of the highest known risk-factors for schizophrenia. This risk quotient is only superseded in those individuals where both biological parents have schizophrenia, or in those with a monozygotic twin also diagnosed with the disorder (Murphy & Owen, 2001) (but see (Mulle, 2015) for evidence of a deletion syndrome with a potentially higher risk for schizophrenia). With a 30-fold increased risk of developing psychosis when compared to the general population (Weisman et al., 2017), about 30% of individuals with 22q11.2DS receive a diagnosis of schizophrenia (Bassett & Chow, 1999; Monks et al., 2014; Murphy, Jones, & Owen, 1999), though lower prevalence has also been reported (Hoeffding et al., 2017). With approximately half of the adolescents with 22q11.2DS showing schizotypical traits and experiencing transient psychotic states (Baker & Skuse, 2005), subthreshold psychotic symptoms appear to present early in this group. Importantly, neither the clinical presentation, nor the clinical path leading to psychosis appear to significantly differ between idiopathic and 22q11.2DS-associated schizophrenia (Hans, Auerbach, Asarnow, Styr, & Marcus, 2000; Welham, Isohanni, Jones, & McGrath, 2009). For instance, among other deficits described across perception and cognition, executive function has been implicated as one of the key domains in the development of psychotic experiences in both syndromic (Chawner et al., 2019; Maeder et al., 2016; Tang & Gur, 2017) and idiopathic schizophrenia (e.g., (Niarchou, Zammit, & Lewis, 2015; Rossi et al., 2016)).

Executive function is an umbrella term used to describe the set of control processes that govern goal-directed behavior and serve to optimize performance on complex cognitive tasks, allowing one to behave flexibly and to adapt to novel, changing circumstances (Gilbert & Burgess, 2008). Executive function abilities—such as working memory, cognitive flexibility, and response inhibition—are critical for academic, professional, and social achievements. To define, plan, and execute daily goals, for example, one must engage working memory to maintain objectives, response inhibition to prevent responses to task-irrelevant distracting information, and cognitive flexibility to adapt to the shifting demands of one’s environment (Baez et al., 2019).

Response inhibition, the process by which one suppresses a prepotent response that might be irrelevant or inappropriate in a given context, is clearly essential for adjusting behavior dynamically with changing environmental contexts (Aron, 2011; Fryer et al., 2018; Hester, Foxe, Molholm, Shpaner, & Garavan, 2005). In 22q11.2DS, behavioral (Maeder et al., 2016; Shapiro, Tassone, Choudhary, & Simon, 2014; Shapiro, Wong, & Simon, 2013; Woodin et al., 2001) and functional magnetic resonance imaging (fMRI) (Gothelf et al., 2007) data point to differences in this domain of executive function. Complicating interpretation, however, these differences are not consistently observed. For example, no behavioral differences in inhibition tasks were found between 22q11.2DS and their siblings in one study (Campbell et al., 2010), or between 22q11.2DS and neurotypical controls in another (Gothelf et al., 2007). Very little is likewise known about response inhibition at prodromal and early stages of schizophrenia, though those individuals seem to show slowed and variable motor responses and decreased engagement of regions implicated in inhibitory control (Fryer et al., 2018). In contrast, behavioral (Badcock, Michie, Johnson, & Combrinck, 2002; Bellgrove et al., 2006; De Sanctis et al., 2012; Enticott, Ogloff, & Bradshaw, 2008; Lipszyc & Schachar, 2010), electrophysiological (EEG) (Chun et al., 2013; Ford et al., 2004; Hughes, Fulham, Johnston, & Michie, 2012; Kiehl, Smith, Hare, & Liddle, 2000; Krakowski et al., 2016; Weisbrod, Kiefer, Marzinzik, & Spitzer, 2000), and fMRI techniques (Ford et al., 2004; Hughes et al., 2012; Kaladjian et al., 2007; Rubia et al., 2001) reveal clear differences in inhibitory processes in chronic schizophrenia.

To gain insight into executive functioning in those at-risk for schizophrenia and better understand the potential of the measures to be used as markers of vulnerability and conversion to psychosis, we assessed response inhibition in individuals with 22q11.2DS—an at-risk for psychosis population—with and without psychotic symptomatology, and compared their behavioral and neural responses to well-matched neurotypical control and idiopathic schizophrenia groups. We further measured the relationship between brain responses and cognitive function. Using standardized cognitive measures and a Go/No-Go EEG task, our goal was to investigate potential markers of risk and disease, dissociating aspects related to 22q11.2DS more broadly from those associated with the presence of psychosis. The analyses focused on already well-characterized cognitive event-related potential (ERP) components that are typically evoked during similar Go/No-Go tasks: The No-Go N2, a negative-going ERP component peaking between 200 and 300ms and representing early, automatic inhibitory (De Sanctis, Butler, Malcolm, & Foxe, 2014; Eimer, 1993; Malcolm, Foxe, Butler, & De Sanctis, 2015; O’Connell et al., 2009) and/or conflict detection processes (Dockree, Kelly, Robertson, Reilly, & Foxe, 2005; Donkers & Van Boxtel, 2004; Morie et al., 2014); the No-Go P3, a positive potential that peaks at about 300-500ms, argued as a marker of response inhibition (Bokura, Yamaguchi, & Kobayashi, 2001; Groom & Cragg, 2015; Kiefer, Marzinzik, Weisbrod, Scherg, & Spitzer, 1998; Waller, Hazeltine, & Wessel, 2019; Wessel & Aron, 2015), stimulus evaluation (Benvenuti, Sarlo, Buodo, Mento, & Palomba, 2015; Bruin & Wijers, 2002; Smith, Johnstone, & Barry, 2008) and adaptive, more effortful forms of control (De Sanctis et al., 2014; Malcolm et al., 2015; Wiersema & Roeyers, 2009); the error-related negativity (ERN or Ne), a component occurring within 100ms of an erroneous response, argued to reflect a mismatch between response selection and response execution (Falkenstein, Hohnsbein, Hoormann, & Blanke, 1991; Nieuwenhuis, Ridderinkhof, Blom, Band, & Kok, 2001), but not remedial action (Nieuwenhuis et al., 2001); and the error-related positivity (Pe), a component peaking between 200 and 500ms post incorrect-response, which has been suggested to reflect conscious error processing or updating of error context (Leuthold & Sommer, 1999; Nieuwenhuis et al., 2001). Additionally, given the visual nature of the Go/No-Go task employed here and the reported differences in early visual-evoked potentials in schizophrenia, when compared to the neurotypical population (Butler & Javitt, 2005; Foxe, Doniger, & Javitt, 2001; Foxe, Murray, & Javitt, 2005; Foxe, Yeap, & Leavitt, 2013; Yeap, Kelly, Sehatpour, et al., 2008) and in 22q11.2DS (Biria et al., 2018; Magnee, Lamme, de Sain-van der Velden, Vorstman, & Kemner, 2011), the early visual components P1, N1, and P2 were also examined. This allowed us to further investigate the relationships between sensory-perceptual and response inhibition neural responses. While those ERP components with potential as markers of risk were expected to be reduced in 22q11.2DS regardless of the presence of psychotic symptoms, components with potential as early neuromarkers of disease were expected to be reduced only in those with 22q11.2DS and psychotic symptomatology.

## MATERIALS AND METHODS

### Participants

Twenty-eight participants diagnosed with 22q11.2DS (22q; age range: 14-35 years old; 14 with at least one psychotic symptom) and 15 individuals diagnosed with schizophrenia (SZ; age range: 18-63 years old) were recruited. Given the age differences between the two groups, two neurotypical control groups were recruited: one age-matched to the 22q11.2DS sample (NT 22q; N=27, age range: 14-38 years old); the other age-matched to the schizophrenia sample (NT SZ; N=15, age range: 25-61 years old). Individuals with 22q11.2DS were recruited via social media and the Montefiore-Einstein Regional Center for 22q11.2 Deletion Syndrome, whereas individuals diagnosed with schizophrenia were recruited through referrals from clinicians in the Department of Psychiatry at Montefiore and Jacobi health systems and through flyers placed at these clinical sites. The recruitment of neurotypical controls was primarily done by contacting individuals from a laboratory-maintained database and through flyers. Exclusionary criteria for the neurotypical groups included hearing impairment, developmental and/or educational difficulties or delays, neurological problems, and the presence of psychotic symptomatology or of any other psychiatric diagnosis. Exclusionary criteria for the 22q11.2DS and the schizophrenia groups included hearing impairment and current neurological problems. All participants passed a hearing screening (thresholds below 25dB NHL for 500, 1000, 2000, and 4000Hz) performed on both ears using a *Beltone Audiometer* (Model 112). One participant in the 22q11.2DS group (with psychotic symptomatology) was unable to perform the EEG task and their data were therefore not included in the analyses. All participants signed an informed consent approved by the Institutional Review Board of the Albert Einstein College of Medicine and were monetarily compensated for their time.

### Experimental Procedure and Stimuli

Testing was carried out over 2 visits and included cognitive testing and EEG recordings. Cognitive testing focused on measures of intelligence and response inhibition. The Wechsler Adult Intelligence Scale, WAIS-IV (Wechsler, 2008) or the Wechsler Intelligence Scale for Children, WISC-V (Wechsler, 2014) were used, depending on the age of the participant. The IQ measure used refers to the Full-Scale IQ index. Two individuals with 22q11.2DS had already been IQ tested using a Wechsler scale within the previous six months and were therefore not retested in-house. To assess response inhibition, the Color-Word Interference Test from the Delis-Kaplan Executive Function System (D-KEFS; Delis, Kaplan, & Kramer, 2001) and the Conners Continuous Performance Test 3 (CPT; (Conners, 2000)) were used. The Color-Word Interference Test consists of four parts: color naming, word reading, inhibition, and inhibition/switching, for which both speed and accuracy (number of errors) are measured. Given the focus of the current study, only the inhibition score is included. The score reported reflects a combined measure of speed and accuracy, computed using the inverse efficiency score (*IES=RT/(1-PE)*, where *RT* is the individual’s average reaction time in the condition, and *PE* is the subject’s proportion of errors in the condition (Townsend & Ashby, 1978). The inhibition score is only available for a subset of the individuals with 22q11.2DS (N=19; the remainder were unable to understand the instructions (N=2), were color blind (N=1) or were part of an initial data collection in which the D-KEFS was not administered). Additionally, six individuals in the NT 22q group and two in the NT SZ group did not complete cognitive testing and thus are not included in the summary IQ statistics and in the correlational analyses. For the CPT, standardized scores of commission errors (incorrect responses to non-targets) and perseverations (random, repetitive, or anticipatory responses), both associated with response inhibition (Lin, Tseng, Lai, Matsuo, & Gau, 2015), are reported. Due to software malfunctioning, CPT measurements are missing for five individuals in the NT 22q group, five in the 22q group, four in the NT SZ group, and one in the SZ group. To test for the presence of psychotic symptomatology (and co-morbidities), either the Structured Clinical Interview for DSM-5, SCID-V (First, Williams, Karg, & Spitzer, 2015) or the Structured Clinical Interview for DSM-IV Childhood Diagnoses, Kid-SCID (Hien et al., 1994) was performed.

During the EEG session, participants performed a Go/No-Go task in which they were asked to respond quickly and accurately to every stimulus presentation, while withholding responses to the second instance of any stimulus repeated twice in a row. The probability of Go and No-Go trials was .85 and .15, respectively. Positively and neutrally valenced pictures from the International Affective Picture System (IAPS; Lang and Cuthbert, 1997), a set of normative photographs depicting people, landscapes, abstract patterns, and objects (http://csea.phhp.ufl.edu/Media.html#topmedia), were presented in a pseudorandom sequence. Stimuli, subtended 8.6° horizontally by 6.5° vertically, were presented centrally every 1000 ms on average for 600 ms with a (random) inter-stimulus-interval between 350 and 450ms. Three 12-minute blocks were run. Each block consisted of 540 trials, for a total of 1620 per participant, 243 of which were inhibition trials.

### Data acquisition and analysis

EEG data were acquired continuously at a sampling rate of 512 Hz from 64 locations using scalp electrodes mounted in an elastic cap (Active 2 system; Biosemi^tm^, The Netherlands; 10-20 montage). Preprocessing was done using EEGLAB (version 14.1.1) (Delorme & Makeig, 2004) toolbox for MATLAB (version 2017a; MathWorks, Natick, MA). Data were downsampled to 256 Hz, re-referenced to the average and filtered using a 0.1 Hz high pass filter (0.1 Hz transition bandwidth, filter order 16896) and a 45 Hz low pass filter (11 Hz transition bandwidth, filter order 152). Both were zero-phase Hamming windowed sinc FIR filters. Bad channels were automatically detected based on kurtosis measures and rejected after visual confirmation. Ocular artifacts were removed by running an Independent Component Analysis (ICA) to exclude components accounting for eye blinks and saccades. After ICA, the previously excluded channels were interpolated, using the spherical spline method. Data were segmented into epochs of −100ms to 1000ms using a baseline of −100ms to 0ms. For the error-related activity analyses, data were segmented into epochs of −100ms to 700ms (response-locked) using a baseline of −100ms to 0ms. These epochs went through an artifact detection algorithm (moving window peak-to-peak threshold at 120 μV). Given that differences were found between the number of trials per group (because some individuals completed two instead of three blocks, although all subjects included had a trial exclusion percentage below 30%, there were still differences between and within groups), to equate number of trials per participant, 200 trials for hits, 50 trials for correct rejections, and 50 trials for false alarms were chosen randomly per subject.

P1 was measured between 90 and 130ms, N1 between 160 and 200ms, and P2 between 230 and 280ms. The amplitudes used for the statistics are an average (across each of the time windows) of the signal at PO7 and PO8 for hits only. N2 was measured between 210 and 240ms and P3 between 350 and 500ms at CPz. For the N2 and the P3, both correct rejections (i.e. successfully withheld responses to a repeated image) and hits (and the difference between the two) were included in the analyses. The error-related negativity (Ne) was measured between 0 and 50ms at FCz. The error-related positivity (Pe) was measured between 200 and 400ms at CPz. These mean amplitude data were used for both between-groups statistics and Spearman correlations. No correlations were performed differentiating the two 22q11.2DS groups due to the small number of individuals per group. Behavioral measures (accuracy and reaction time) were additionally taken during the EEG task. Hits were defined as responses to a non-repeated picture; correct rejections as the absence of response to a repeated picture; false alarms as responses to a repeated picture. Only hits and correct rejections preceded by a hit were included. *D*-prime (d’ = z(H) - z(F)) was calculated per subject. All *p-values* (from *t*-tests and Spearman correlations) were submitted to Holm-Bonferroni corrections for multiple comparisons (Holm, 1979), using the *p.adjust* of the *stats* package in R (R Core Team, 2014).

Two levels of analyses were carried out. First, the 22q11.2DS and schizophrenia groups were compared to their respective age-matched control groups. Mixed-effects models were implemented separately per group (NT 22q *versus* 22q; NT SZ *versus* SZ) to analyze trial-by-trial data, using the *lmer* function in the *lme4* package (D. Bates, Mächler, Bolker, & Walker, 2014) in R (RCoreTeam, 2014). Group was always a fixed factor, and trial type an additional numeric fixed factor for reaction times and P3 analyses. Subjects and trials were added as random factors. Models were fit using the maximum likelihood criterion. *P* values were estimated using *Satterthwaite* approximations. Second, given that half of the individuals with 22q11.2DS tested here presented at least one psychotic symptom (N=14; 22q11.2DS+), we divided the 22q11.2DS group into two sub-groups (22q11.2DS+ and 22q11.2DS-, N=13) and compared them to age-matched neurotypical controls (NT 22q) and to the schizophrenia group. Mixed-effects models were implemented as above, but age was now included in the models to control for the variance potentially explained by the age differences between the groups (with the schizophrenia sample being significantly older than the NT 22q and the two 22q11.2DS groups).

## RESULTS

### Demographics and cognitive function measures

Table 1 shows a summary of the included participants’ age and biological sex. Two-sample independent-means *t* tests were run in R (RCoreTeam, 2014) to test for group differences. In cases in which the assumption of the homogeneity of variances was violated, *Welch* corrections were applied to adjust the degrees of freedom. There were more females than males in the NT 22q and 22q groups and more males than females in the NT SZ and SZ groups, but no differences in biological sex between groups. Likewise, age did not differ between the NT 22q and 22q groups and the NT SZ and SZ groups. No differences in sex or age were found between the 22q11.2DS- and the 22q11.2DS+ groups. 25.9% of those with 22q11.2DS were diagnosed with a mood disorder, 25.9% with an anxiety disorder, and 7.4% with schizophrenia. From those with schizophrenia, 6.7% met criteria for a mood disorder and 13.3% for an anxiety disorder. 33.3% of those with 22q11.2DS were taking antidepressants, 25.9% anticonvulsants, 11.1% antipsychotics, 7.4% antimanics, and 7.4% stimulants. From those individuals diagnosed with schizophrenia, 86.7% were taking antipsychotics, 33.3% anticholinergics, 26.7% anticonvulsants, and 13.3% antidepressants. One individual with schizophrenia was taking testosterone.

**Table 1.** Characterization of the neurotypical, 22q11.2DS (overall and per sub-group: 22q11.2DS- and 22q11.2DS+), and schizophrenia groups included in the analyses: age and biological sex.

|  | age | biological sex |
| --- | --- | --- |
| **NT 22q** | M=21.92; SD=6.70 | 11 males, 16 females |
| **22q** | M=21.89; SD=6.66 | 11 males, 16 females |
| t-test | $t=0.02, df=51.99, p=.98$ | - |
| Cohen's d | $d=0.01$ | - |
| **22q-** | M=22.46; SD=7.34 | 7 males, 6 females |
| **22q+** | M=21.36; SD=6.18 | 4 males, 10 females |
| t-test | $t=0.42, df=23.57, p=.68$ | - |
| Cohen's d | $d=0.16$ | - |
| **NT SZ** | M=43.93; SD=10.33 | 9 males, 6 females |
| **SZ** | M=41.20; SD=11.66 | 9 males, 6 females |
| t-test | $t=0.68, df=27.59, p=.50$ | - |
| Cohen's d | $d=0.25$ | - |

Figure 1 and table 2 show a summary of the included participants’ performance on the IQ, D-KEFS, and CPT measures.

**Figure 1.**
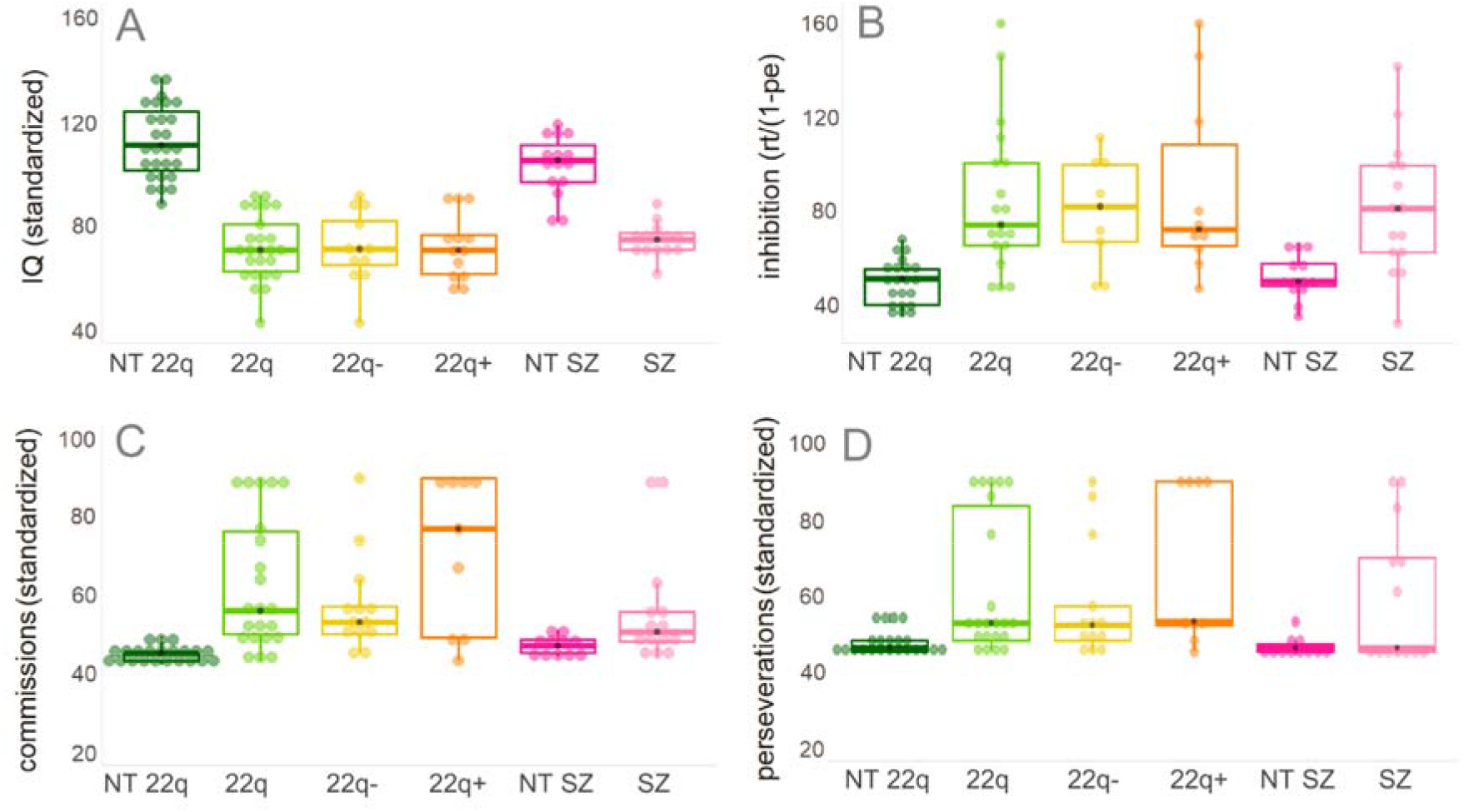
Included participants’ performance on IQ (panel A), D-KEFS (panel B), and CPT (panels C and D) measures. In panels B, C, and D, higher scores represent worse scores.

**Table 2.** Characterization of the neurotypical, 22q11.2DS (overall and per sub-group: 22q11.2DS- and 22q11.2DS+), and schizophrenia groups included in the analyses: cognitive measures

|  | **IQ** | **inhibition (D-KEFS)** | **commissions (CPT)** | **perseverations (CPT)** |
| --- | --- | --- | --- | --- |
| **NT 22q** | M=111.93; SD=14.20 | M=49.49; SD=9.63 | M=44.73; SD=1.96 | M=47.36; SD=3.22 |
| **22q** | M=71.16; SD=12.96 | M=84.27; SD=31.94 | M=63.00; SD=17.13 | M=61.77; SD=18.46 |
| t-test | $t=10.82, df=49.99, p=.01$ | $t=-4.56, df=20.96, p=.01$ | $t=-4.97, df=21.55, p=.01$ | $t=-3.62, df=22.28, p=.01$ |
| Cohen's d | $d=3.00$ | $d=-1.47$ | $d=-1.50$ | $d=-1.09$ |
| **22q-** | M=71.08; SD=13.98 | M=79.53; SD=22.93 | M=57.23; SD=12.52 | M=57.54; SD=15.74 |
| **22q+** | M=71.23; SD=12.52 | M=88.54; SD=39.13 | M=71.33; SD=20.06 | M=67.89; SD=21.13 |
| t-test | $t=-0.03, df=22.18, p=.98$ | $t=-0.62, df=14.76, p=.54$ | $t=-1.87, df=12.30, p=.08$ | $t=-1.25, df=13.96, p=.23$ |
| Cohen's d | $d=0.01$ | $d=0.28$ | $d=0.84$ | $d=0.54$ |
| **NT SZ** | M=103.20; SD=11.83 | M=51.63; SD=9.26 | M=46.91; SD=2.47 | M=46.54; SD=2.46 |
| **SZ** | M=74.00; SD=6.41 | M=81.49; SD=28.72 | M=56.14; SD=14.76 | M=59.77; SD=18.42 |
| t-test | $t=8.34, df=21.86, p=.01$ | $t=-3.80, df=17.27, p=.01$ | $t=-2.30, df=13.92, p=.04$ | $t=-2.56, df=12.51, p=.04$ |
| Cohen's d | $d=3.07$ | $d=-1.40$ | $d=-0.87$ | $d=-1.01$ |

Statistical analyses, conducted as described above, confirmed that the groups differed significantly in IQ, and in the D-KEFS and CPT measures, with both 22q and SZ groups performing worse than their neurotypical peers (Table 2). No differences were found between the 22q11.2DS- and the 22q11.2DS+ groups.

### Go/No-Go EEG task

#### Behavioral performance

Figure 2 and Table 3 show the participants’ behavioral performance (*d*-prime and reaction times) on the Go/No-Go EEG task. To test for differences in *d*-prime between the groups, two-sample independent-means *t* tests were run in R (RCoreTeam, 2014) as described above. When compared to their neurotypical controls, individuals diagnosed with 22q11.2DS (*t*=3.45, *df*=48.17, *p*=.01, *d*=0.95) and those with schizophrenia (*t*=2.27, *df*=25.53, *p*=.03, *d*=0.83) presented lower *d*-prime scores, reflecting lower rates of hits in both clinical populations when compared to the control groups (NT-22q: *t*=3.68, *df*=27.24, *p*=.01, *d*=1.01; NT-SZ: *t*=2.85, *df*=16.31, *p*=.04, *d*=1.04). To test for differences between the groups in reaction time, mixed-effects models were implemented as described above. Overall effects of group were found: Individuals with 22q11.2DS (*ß* = 37.74, SE = 14.48, *p* = .01) and those with schizophrenia (*ß* = 104.85, SE = 31.82, *p* = .01) responded slower than their neurotypical peers. An effect of trial type was also observed, with false alarms (NT-22q: *ß* = −35.61, SE = 2.92, *p* = .01; NT-SZ: *ß* = −33.49, SE = 4.48, *p* = .01) and hits after false alarms (NT-22q: *ß* = −49.78, SE = 3.08, *p* = .01; NT-SZ: *ß* = −38.22, SE = 4.72, *p* = .01) resulting in shorter reaction times than hits. Lastly, the interactions between group and trial type were significant. Differently from what was observed in the neurotypical control group, in 22q11.2DS, reaction times increased for false alarms when compared to hits (*ß* = 41.03, SE = 3.81, *p* = .01) and for hits after false alarms when compared to hits (*ß* = 86.83, SE = 4.04, *p* = .01). A similar pattern was found in schizophrenia, with longer reaction times in hits after false alarms when compared to hits (*ß* = 61.36, SE = 6.27, *p* = .01) (see Figure 2, panels B to D).

**Figure 2.**
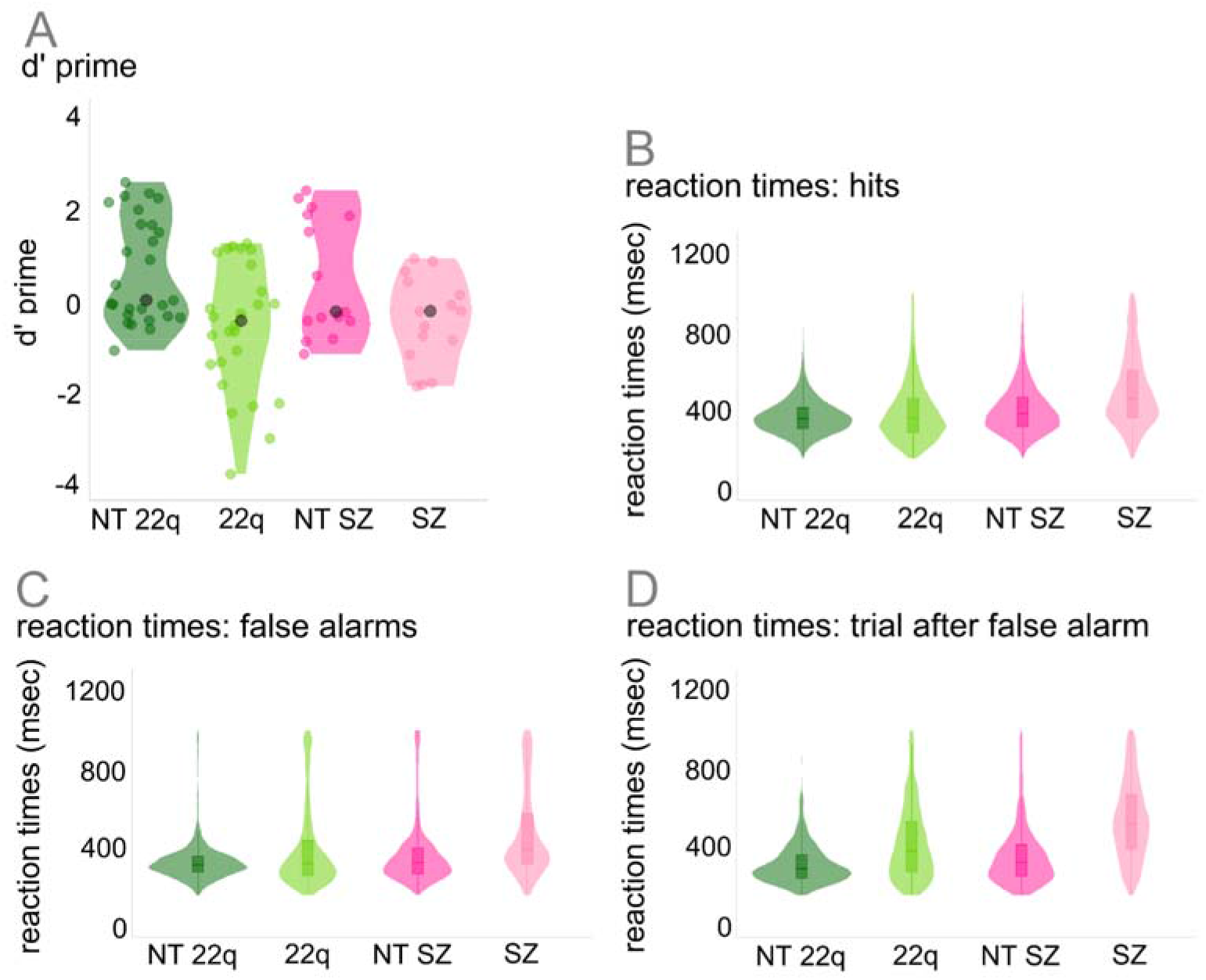
Participants’ behavioral performance on the Go/No-Go EEG task.

**Table 3.** Participants’ behavioral performance on the Go/No-Go EEG task: D-prime and reaction times

|  | d-prime | RT hits | RT false alarms | RT after false alarm |
| --- | --- | --- | --- | --- |
| NT 22q | M=0.62; SD=1.15 | M=366.19; SD=90.48 | M=324.16; SD=103.74 | M=311.06; SD=101.76 |
| 22q | M=-0.61; SD=1.42 | M=389.43; SD=150.46 | M=388.23; SD=208.31 | M=419.48; SD=190.20 |
| 22q- | M=-0.54; SD=1.35 | M=379.22; SD=132.63 | M=370.85; SD=190.42 | M=422.55; SD=188.93 |
| 22q+ | M=-0.67; SD=1.53 | M=400.25; SD=166.62 | M=403.29; SD=221.64 | M=416.87; SD=191.32 |
| NT SZ | M=0.50; SD=1.32 | M=404.12; SD=128.83 | M=363.48; SD=168.05 | M=356.86; SD=156.61 |
| SZ | M=-0.46; SD=0.95 | M=497.36; SD=184.06 | M=468.36; SD=220.79 | M=536.09; SD=198.11 |

As can be appreciated in Figure 3 (panel A) and Table 3, both 22q11.2DS groups performed worse than the controls in terms of accuracy (overall effect of group: *F*(2,50)=5.92, *p* = .01, η^2^ = 0.19; post-hoc comparisons with neurotypical controls, holm-corrected: 22q112.DS-: *p* = .021, 22q112.DS+: *p* = .02), but no differences were found between the 22q11.2DS groups and the schizophrenia sample (*F*(2,35)=0.10, *p* = .90, η^2^ = 0.01). To control for the difference in age between the 22q11.2DS and the schizophrenia group, age was added to the ANOVA model. There were no effects of age (*F*(1,35)=1.16, *p* = .29, η^2^ = 0.03) or of the interaction between group and age (*F*(2,35)=3.17, *p* = .06, η^2^ = 0.15). Figure 3 (panel B) further indicates that both 22q11.2DS- and 22q11.2DS+ groups were faster compared to the schizophrenia group (22q11.2DS-: *ß* = −122.18, SE = 28.44, *p* = .01; 22q11.2DS+: *ß* = −84.54, SE = 27.89, *p* = .01). When compared to the neurotypical controls, the 22q11.2DS-group was as fast (*ß* = 25.60, SE = 17.33, *p* = .14), whereas the 22q11.2DS+ group was slower (*ß* = 62.89, SE = 16.91, *p* = .01).

**Figure 3.**
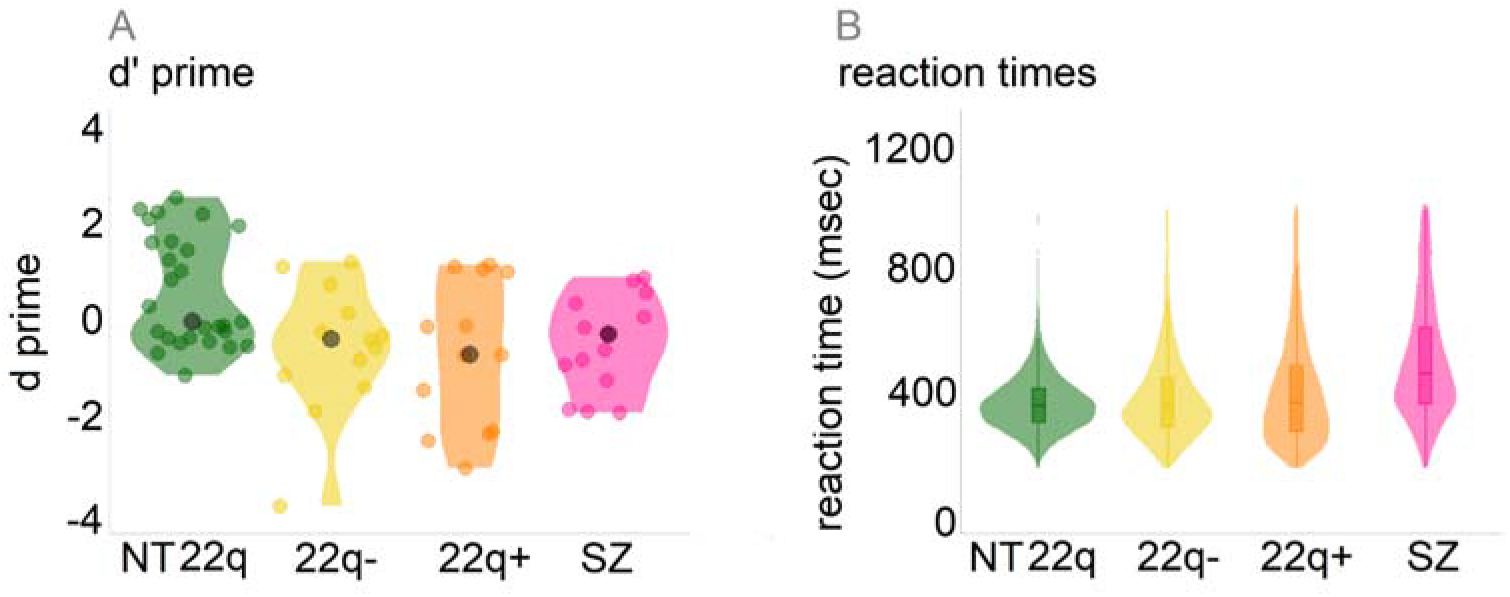
Participants’ behavioral performance on the Go/No-Go EEG task (22q11.2DS- and 22q11.2DS+ groups compared to the neurotypical control (NT 22q) and schizophrenia groups).

#### Basic visual processing: P1, N1, and P2

Figure 4 shows the averaged ERPs and topographies for the early sensory time windows of interest (P1, N1, and P2), per channel (Figure 3A) and by group (see supplementary Figure 1 for additional scalp locations). Only hits are included in these analyses. Mixed-effects models were implemented as described in the Methods Section. Amplitude (PO7-PO8 average) at each trial was the numeric dependent variable. Though both clinical groups’ data appeared to be characterized by slightly decreased visual evoked responses (Figure 3A), neither 22q11.2DS (P1: *ß* = −1.72, SE = 1.23, *p* = .17; N1: *ß* = −1.70, SE = 1.42, *p* = .24; P2: *ß* = −2.62, SE = 1.50, *p* = .08) nor schizophrenia (P1: *ß* = −1.28, SE = 1.17, *p* = .28; N1: *ß* = - 0.75, SE = 1.30, *p* = .57; P2: *ß* = −2.80, SE = 1.69, *p* = .11) groups differed significantly from their controls. To more thoroughly explore the spatio-temporal dynamics of these responses, post-hoc statistical cluster plots were computed (supplementary Figure 2). These plots suggest the existence of differences between individuals with schizophrenia and their controls in parietal channels around the P1 time window.

**Figure 4.**
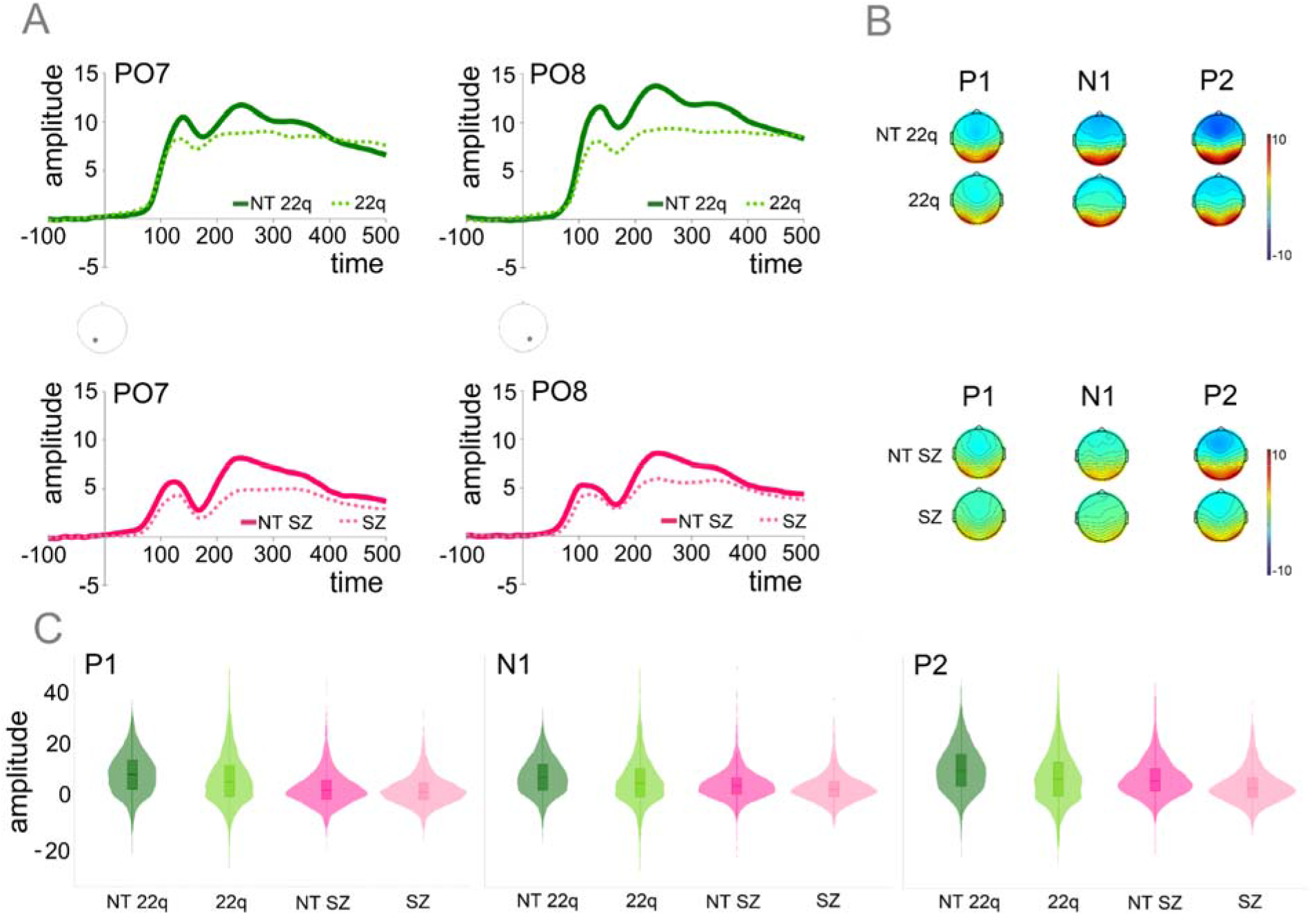
Panel A: Averaged ERPs for the early sensory time windows of interest (P1, N1, and P2), from channels over left (PO7) and right (PO8) posterior scalp regions and by group; Panel B: Topographies for the P1 (90-130 ms), N1 (160-200 ms), and P2 (230-280 ms) time windows, per group; Panel C: Plots showing distribution of amplitudes for P1, N1, and P2 per group at PO7 and PO8 (trial-by-trial data).

Figure 5 shows the averaged ERPs for the early sensory time windows of interest per channel and by 22q11.2DS group (22q11.2DS-*versus* 22q11.2DS+), when compared to neurotypical control and schizophrenia groups. Mixed-effects models were implemented as above, but age was now included in the models. The 22q11.2DS-group did not differ from the neurotypical controls in the P1 (*ß* = −0.39, SE = 1.44, *p* =.78), N1 (*ß* = −1.14, SE = 1.78, *p* = .52), or P2 (*ß* = −1.39, SE = 1.81, *p* =.45) time windows. The 22q11.2DS+ group showed significantly decreased amplitudes when compared to the control group in the P1 (*ß* = −3.29, SE = 1.41, *p* = .02) and in the P2 (*ß* = −4.36, SE = 1.77, *p* = .02), but not in the N1 time window (*ß* = −2.24, SE = 1.73, *p* = .20). The 22q11.2DS+ group did not differ from the schizophrenia group in any of the time windows of interest (P1: *ß* = 0.20, SE = 1.71, *p* =.91; N1: *ß* = 2.20, SE = 1.97, *p* =.27; P2: *ß* = 0.50, SE = 2.04, *p* =.80). Inversely, the 22q11.2DS-group presented increased amplitudes when compared to schizophrenia in P1 (*ß* = 3.88, SE = 1.72, *p* = .03) and P2 (*ß* = 4.74, SE = 2.05, *p* = .02) (but not in N1 (*ß* = 3.59, SE = 1.98, *p* = .08).

**Figure 5.**
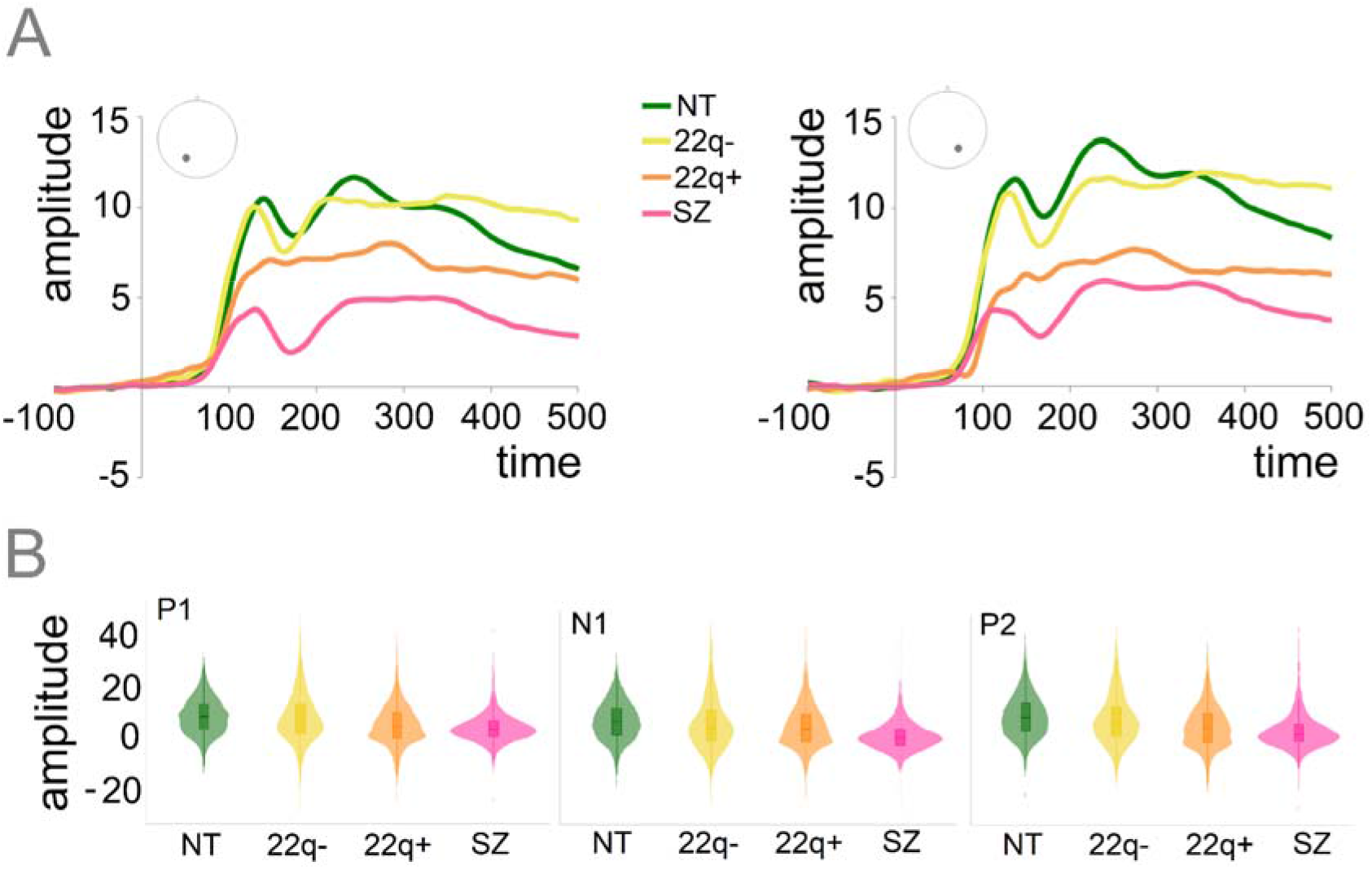
Panel A: Averaged ERPs for the early sensory time windows of interest (P1, N1, and P2), per channel (PO7 and PO8) and by group (NT 22q, 22q+, 22q-, and SZ); Panel B: Plots showing distribution of amplitudes for P1, N1, and P2 per group at PO7 and PO8 (trial-by-trial data).

#### Response inhibition: N2 & P3

Figure 6 shows the averaged ERPs and topographies for N2 and P3 by group and trial type (hits and correct rejections). Mixed-effects models were implemented as described in the Methods Section. In the N2 time window, no significant effects were found for the individuals with 22q11.2DS and their age-matched peers. Likewise, no significant effect of group was found for schizophrenia, though a significant interaction was observed between group and trial type: When compared to their control group, individuals with schizophrenia presented a reduced difference between correct rejections and hits (*ß* = 1.26, SE = 0.46, *p* = .01). A significant effect of trial type was also present, with correct rejections evoking a slightly enlarged N2, but only in the NT-SZ (*ß* = −0.75, SE = 0.37, *p* = .04), not in the NT-22q (*ß* = 0.06, SE = 0.24, *p* = .82). In the P3 time window, the correct rejection versus hit difference was smaller in each of the clinical groups when compared to their controls (Figure 5A), indicating smaller P3s. Though no significant effect of group was found, significant interactions were observed between group and trial type. Here, for both 22q11.2DS (*ß* = −1.11, SE = 0.34, *p* = .01) and schizophrenia (*ß* = −1.56, SE = 0.31, *p* = .01) groups, and when compared to their respective control groups, the difference between correct rejections and hits were, indeed, reduced. A significant effect of trial type was also present, with correct rejections evoking an enlarged P3 (NT-22q: *ß* = 3.18, SE = 0.25, *p* = .01; NT-SZ: *ß* = 2.03, SE = 0.31, *p* = .01).

**Figure 6.**
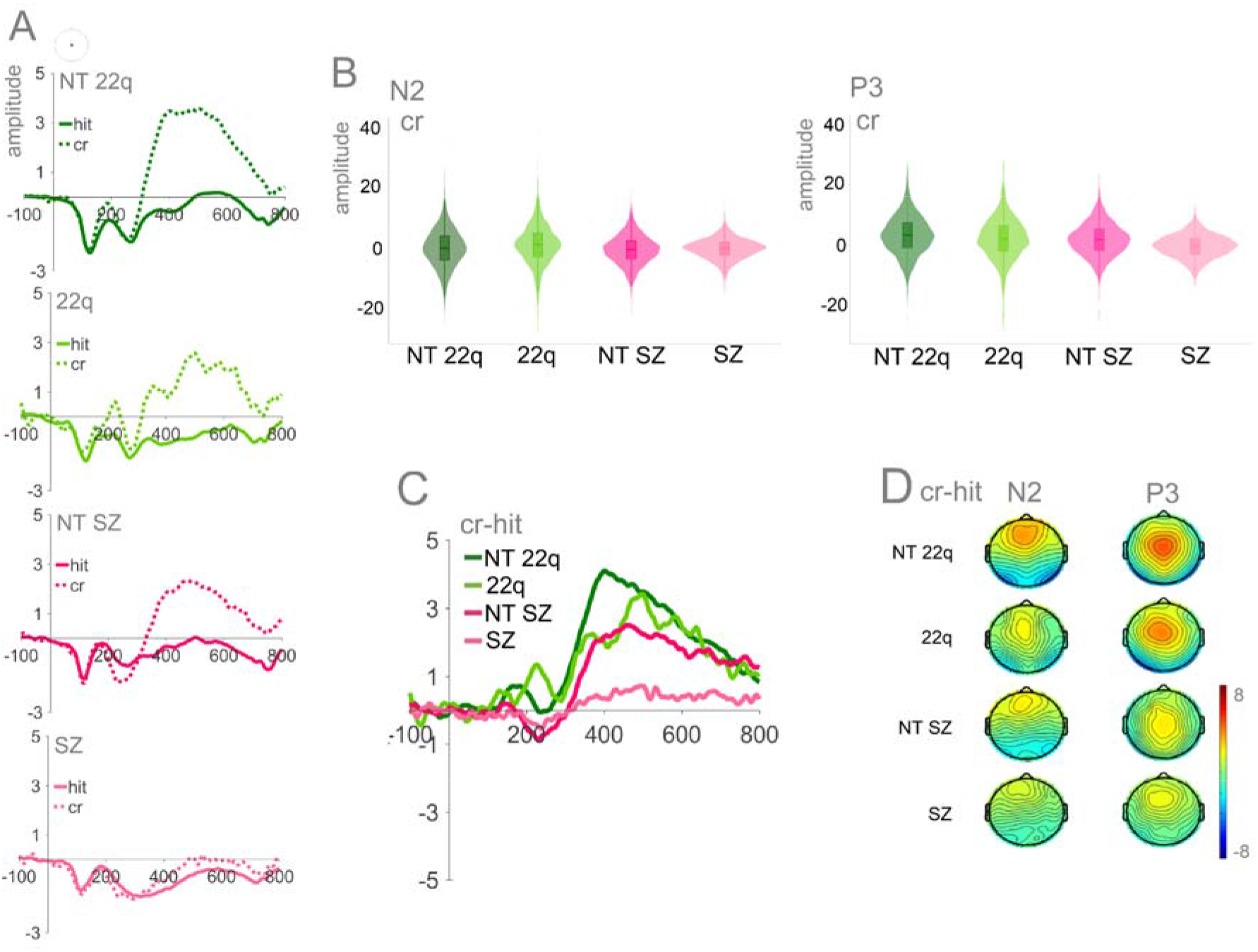
Panel A: Averaged ERPs per group at CPz; Panel B: Plots showing distribution of amplitudes for correct rejections per group at CPz (trial-by-trial data) for N2 and P3. Panel C: Difference waves (correct rejections – hits) per group. Panel D: Topographies for N2 (210-240ms) and P3 (350-500 ms).

Figure 7 depicts the N2/P3 difference waves (correct rejections – hits) for the 22q11.2DS groups (22q11.2DS- *versus* 22q11.2DS+), their neurotypical control group, and the schizophrenia group. Mixed-effects models were implemented as above, except with age now included in the models. In the N2 time window, the only significant difference found was between the 22q11.2DS-group and the age-matched neurotypical controls (*ß* = 1.41, SE = 0.41, *p* = .01), with the former showing an increased difference wave. In the P3 time window, only the 22q11.2DS+ group differed from the neurotypical control group, showing a decreased difference between correct rejections and hits (*ß* = −1.68, SE = 0.42, *p* = .01), while only the 22q11.2DS- group differed significantly from the schizophrenia group, showing an increased difference between correct rejections and hits (*ß* = 3.38, SE = 0.85, *p* = .01).

**Figure 7.**
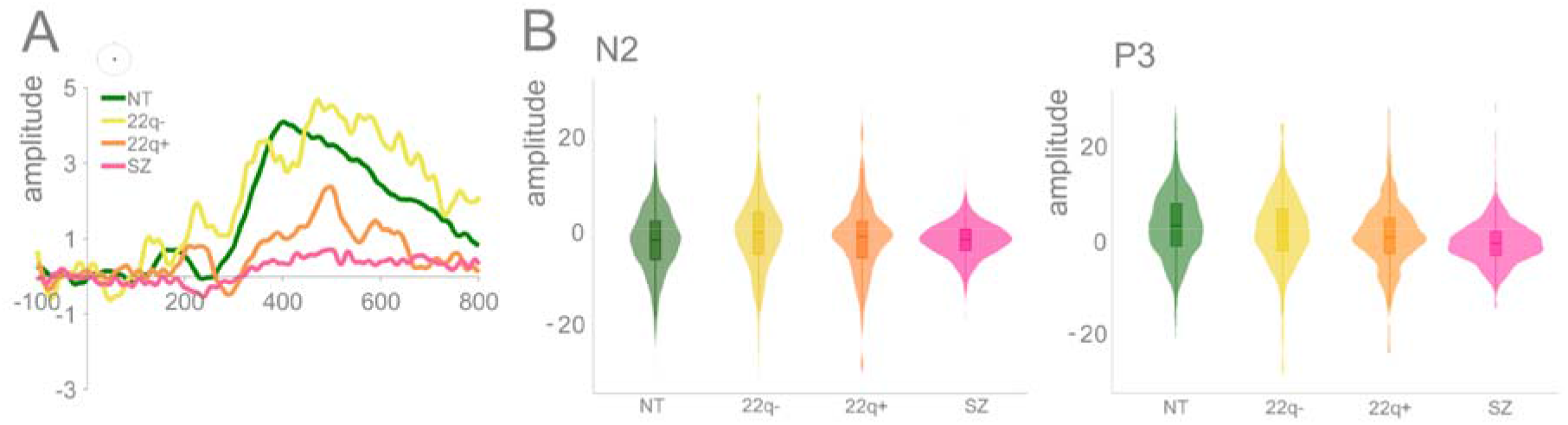
Panel A: N2/P3 (difference waves: correct rejections – hits) per group (NT, 22q-, 22q+, SZ) at CPz. Panel B: Plots showing distribution of amplitudes for correct rejections per group at CPz (trial-by-trial data) for N2 and P3.

#### Error-related activity: Ne & Pe

Figure 8 shows the averaged ERPs and topographies for Ne by group. Mixed-effects models, implemented as described in the Methods Section, revealed that when compared to the neurotypical controls, both those with 22q11.2DS (*ß* = 0.82, SE = 0.36, *p* = .03) and those with schizophrenia (*ß* = 0.97, SE = 0.44, *p* = .04) presented significantly reduced Ne.

**Figure 8.**
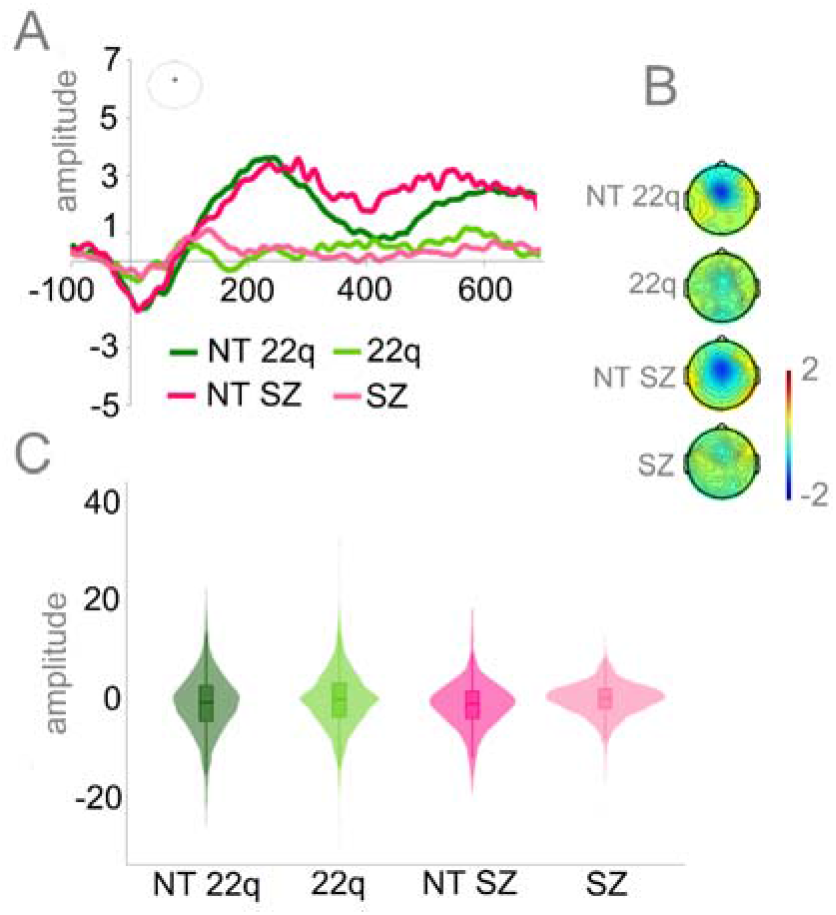
Panel A: Averaged ERPs depicting error-related negativity (Ne) by group at FCz. Panel B: Topographies for the Ne (0-50 ms) per group. Panel C: Plots showing distribution of amplitudes for Ne per group (trial-by-trial data).

Figure 9 shows the averaged ERPs and topographies for Pe by group. Marked differences between each of the clinical groups and their respective control group can be observed (Figure 8A). Mixed-effects models confirmed these observations: When compared to the neurotypical controls, both the individuals diagnosed with 22q11.2DS (*ß* = −3.42, SE = 0.89, *p* < .001) and those with schizophrenia (*ß* = −1.89, SE = 0.73, *p* < .05) presented significantly reduced error-related positivity.

**Figure 9.**
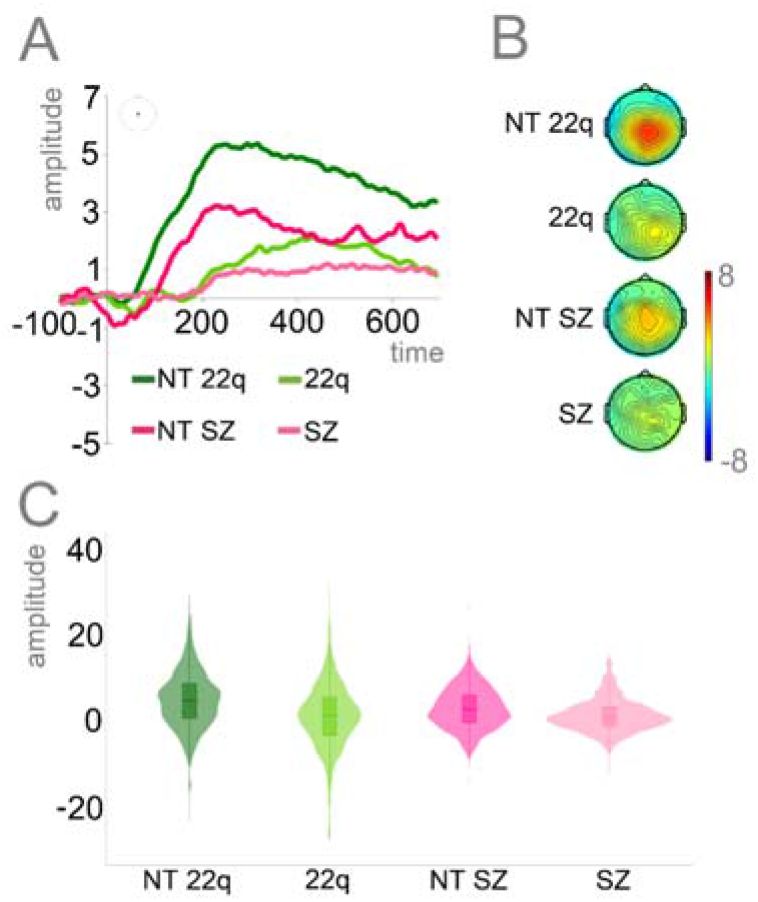
Panel A: Averaged ERPs depicting error-related positivity (Pe) by group at CPz. Panel B: Topographies for the Pe (200-400 ms) per group. Panel C: Plots showing distribution of amplitudes for Pe per group (trial-by-trial data).

Figures 10 and 11 show the Ne and the Pe, respectively, for both of the 22q11.2DS groups (22q11.2DS- *and* 22q11.2DS+) and the neurotypical control and schizophrenia groups. Mixed-effects models were implemented as above, apart from the addition of age to the models. In the Ne time window, while the 22q11.2DS+ group presented reduced amplitudes (*ß* = 1.62, SE = 0.46, *p* = .01), no differences were found between the 22q11.2DS- and the neurotypical controls (*ß* = 0.59, SE = 0.47, *p* = .21) and between either of the 22q11.2DS groups and the schizophrenia group (22q11.2DS-: *ß* = −0.50, SE = 0.34, *p* = .15; 22q11.2DS+: *ß* = 0.65, SE = 0.33, *p* = .06).

**Figure 10.**
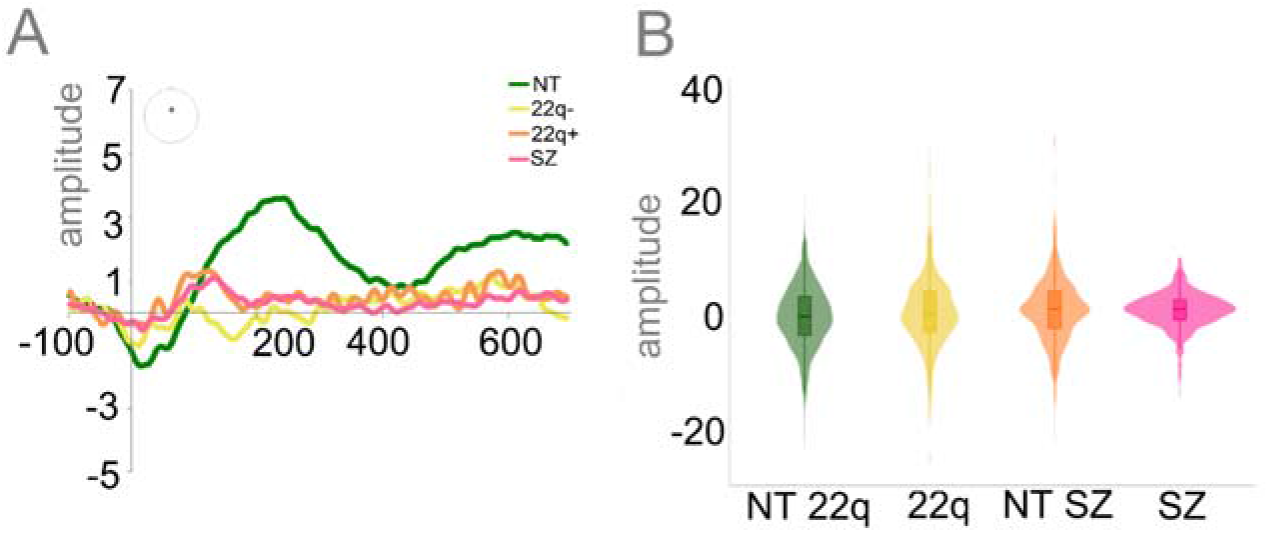
Panel A: Averaged ERPs depicting error-related negativity (Ne) per group (NT, 22q-, 22q+, SZ) at FCz. Panel B: Plots showing distribution of amplitudes for Ne per group (trial-by-trial data).

**Figure 11.**
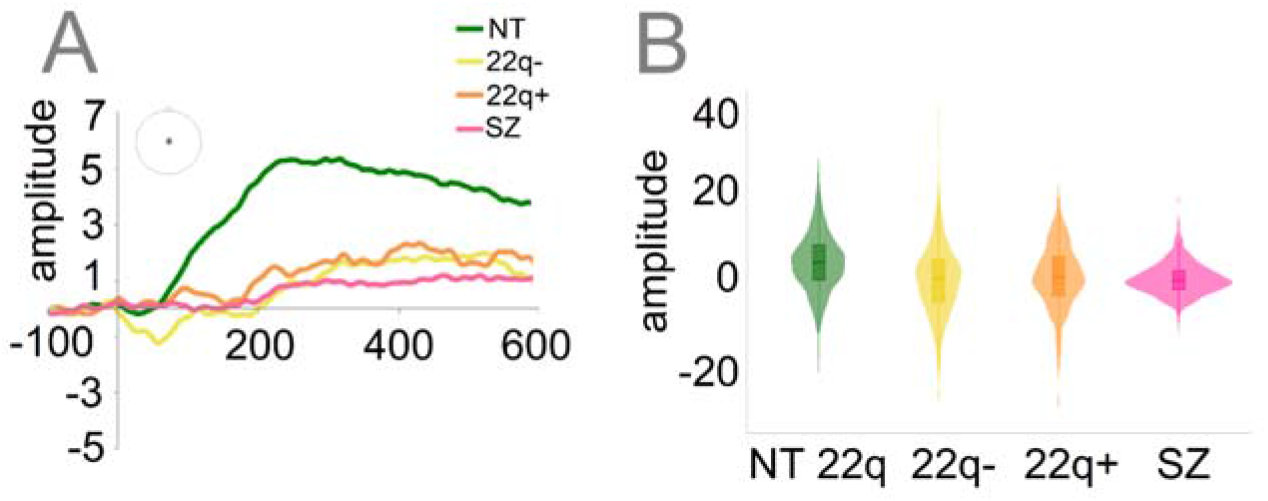
Panel A: Averaged ERPs depicting error-related positivity (Pe) per group (NT, 22q-, 22q+, SZ) at CPz. Panel B: Plots showing distribution of amplitudes for Pe per group (trial-by-trial data).

Differently, both 22q11.2DS groups presented significantly decreased Pe amplitudes when compared to the neurotypical control group (22q11.2DS-: *ß* = −3.44, SE = 1.12, *p* = .01; 22q11.2DS+: *ß* = −2.40, SE = 1.10, *p* = .03), and similar Pe amplitudes when compared to the schizophrenia group (22q11.2DS-: *ß* = 0.51, SE = 1.04, *p* =.63; 22q11.2DS+: *ß* = 0.05, SE = 1.02, *p* = .96).

#### Correlations

Figure 12 shows the significant Spearman correlations between neural responses and clinical scores. P3 amplitude correlated negatively with the inhibition score from the D-KEFS (*r_s_*=-.35, *p* = .04): the smaller the P3 (difference between hits and correct rejections), the higher the inhibition score (with higher scores representing poorer performances) (Figure 12A). Ne amplitudes correlated positively with inhibition (*r_s_*=.39, *p* = .02) and commission errors (*r_s_*=.34, *p* = .04) (Figure 12B). Interestingly, the correlation with inhibition was only significant for the clinical populations, *r_s_*=.52, *p* = .03), not for the neurotypical controls (*r_s_*=-.03, *p* = .87) (Figure 13A). Similarly, Pe amplitudes correlated negatively with inhibition (*r_s_*=-.55, *p* = .02) and commission errors (*r_s_*=-.54, *p* = .02) (Figure 12C). As for the Ne, the correlation between Pe amplitude and inhibition was dictated by the clinical populations (*r_s_*=-.42, *p* = .04), not by the neurotypical controls (*r_s_*=-.16, *p* = .36) (Figure 13B). No correlations were found between N2 and any of the clinical scores.

**Figure 12.**
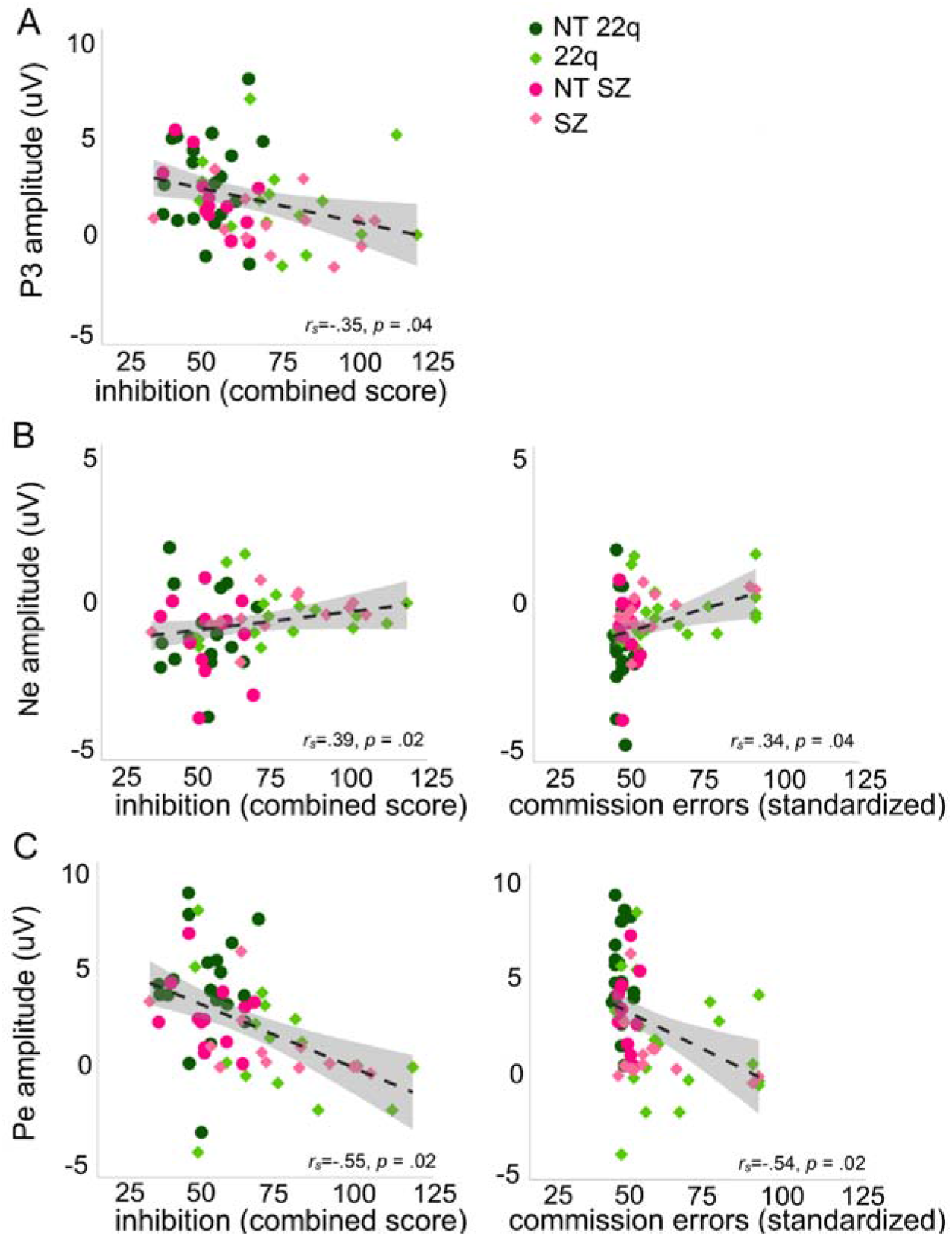
Spearman correlations between P3 amplitude and inhibition (panel A), Ne amplitude and inhibition and commission errors (panel B), Pe amplitude and inhibition and commission errors (panel C).

**Figure 13.**
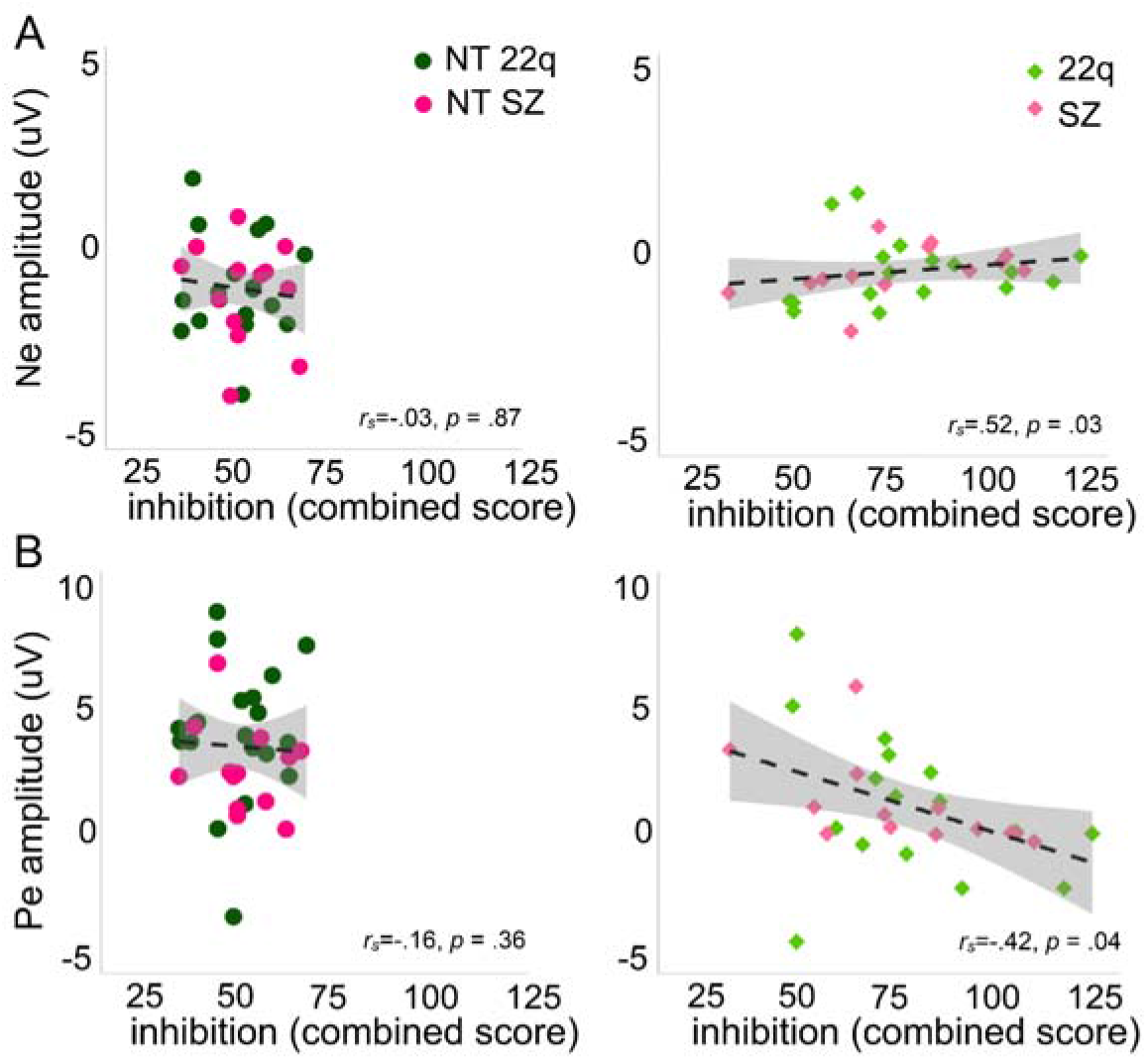
Spearman correlations between Ne amplitude and inhibition per group (neurotypical controls and clinical populations (panel A), and between Pe amplitude and inhibition per group (neurotypical controls and clinical populations (panel B).

Figure 14 depicts the Spearman correlations between the early visual components (P1, N1, and P2) and the response inhibition components N2, P3, Ne, and Pe. P1 correlated positively with P3 (*r_s_*=.42, *p* = .01) and Pe (*r_s_*=.38, *p* = .01), and so did N1 (P3: *r_s_*=.37, *p* = .01; Pe: *r_s_*=.42, *p* = .01) and P2 (P3: *r_s_*=.40, *p* = .01; Pe: *r_s_*=.50, *p* = .01). After correction for multiple comparisons, no significant correlations were found between the early visual components and either N2 or Ne.

**Figure 14.**
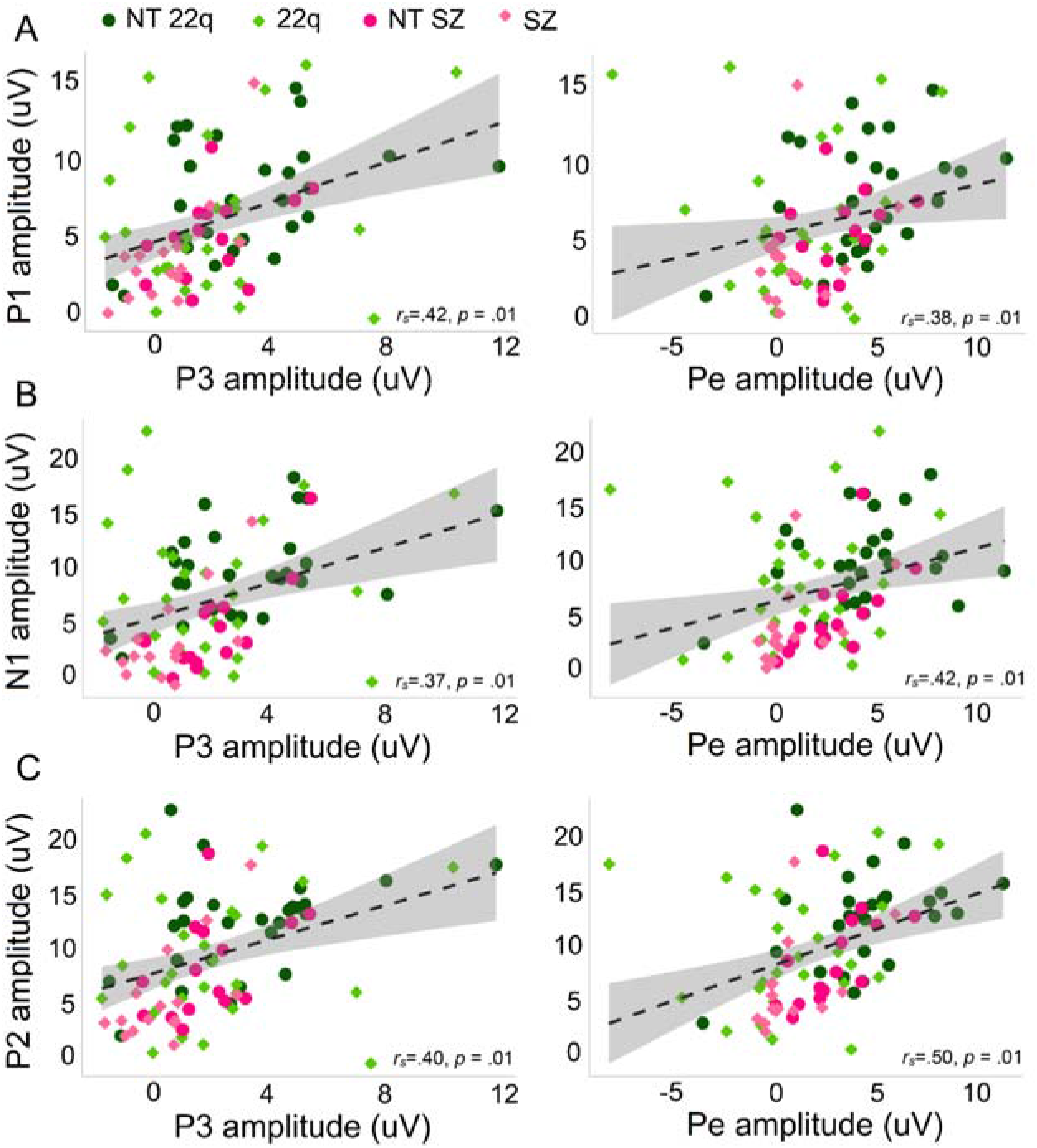
Spearman correlations between early visual components and response inhibition components. Panel A: P1 amplitude and P3 and Pe amplitudes; panel B: N1 amplitude and P3 and Pe amplitudes; panel C: P2 amplitude and P3 and Pe amplitudes

## DISCUSSION

The ability to inhibit inappropriate responses is vital to self-regulation of behavior and cognition, allowing one to adapt planning and actions based on the exigencies of an ever-changing environment. Using standardized cognitive measures and high-density EEG recordings, we characterized the behavioral performance and neural dynamics of response inhibition in individuals with 22q11.2DS, making a specific distinction between those with (22q11.2DS+) and without (22q11.2DS-) psychotic symptomatology. We compared these groups to a cohort of individuals with a frank diagnosis of schizophrenia and to age-matched neurotypical controls, testing for associations between neural and clinical measures of response inhibition. In light of extensive prior reports of early visual processing deficits in schizophrenia (e.g., (Andrade, Butler, Peters, Molholm, & Foxe, 2016)), we also examined the early sensory-perceptual phases of processing in 22q11.2DS to see if this effect would recapitulate in this population.

Neurocognitive assessment showed that individuals with 22q11.2DS and those with schizophrenia, when compared to their neurotypical peers, had greater difficulty inhibiting dominant and automatic verbal (D-KEFS) and motor (CPT) responses. This is in accordance with previous behavioral evidence indicating atypical response inhibition in schizophrenia (Hepp, Maier, Hermle, & Spitzer, 1996) and impaired development of response inhibition in 22q11.2DS (Shapiro et al., 2014). Contrary to what one might expect in light of the cognitive decline that is typically seen to precede psychosis in schizophrenia (Jalbrzikowski & Bearden, 2011; Pantelis et al., 2003), no differences were found here in these measures, or in IQ, in 22q11.2DS individuals with versus without psychotic symptomatology. Arguably, since 22q11.2DS is often characterized by marked cognitive deficits (McDonald-McGinn et al., 2015), one might expect a less noticeable decline preceding psychosis in this population. However, examination of Figure 1 suggests that at least in some of the standardized measures of attention (particularly in inhibition and commission errors), 22q11.2DS- and 22q11.2DS+ performed differently. The lack of detectable significant differences may be driven by the relatively small samples and the increased variability, particularly observed in 22q11.2DS+. And, indeed, a large prospective longitudinal study has demonstrated such differences, where those with 22q11.2DS who developed a psychotic disorder showed a mild cognitive decline relative to those that had not developed a psychotic disorder, particularly in verbal IQ (Vorstman et al., 2015). Here, we did not restrict our sample to those who had a frank psychotic disorder. Instead, we included in the 22q11.2DS+ group any individual who presented psychotic symptomatology, which could further explain the differences between our and Vorstman et al.’s findings.

During the EEG response inhibition task, consistent with what was observed in the D-KEFS and CPT measures, both the 22q11.2DS and the schizophrenia groups displayed poorer *d*-prime scores and longer reaction times compared to their neurotypical peers. In schizophrenia, lower *d*-prime scores (Groom et al., 2008; Simmonite, Bates, Groom, Hollis, & Liddle, 2015) and longer reaction times (Birkett et al., 2007; Krakowski et al., 2016) have been previously reported in the context of Go/No-Go tasks. While lower d-prime scores suggest diminished ability to discriminate between signal and noise, or in this context, to monitor performance in order to adaptively balance demands to detect and rapidly respond or inhibit responses, longer reaction times may be a consequence of the motor slowing described in this population (Morrens, Hulstijn, & Sabbe, 2007). Poorer behavioral performance has likewise been reported in 22q11.2DS (Jonas et al., 2015), but not consistently so. For example, in an fMRI study investigating response inhibition, adolescents with 22q11.2DS did not differ from their age-matched peers in hit and false alarm rates and reaction times, despite the presence of between-groups differences in activation in a region of the left parietal lobe. The authors speculated that this was due to the ability of those with 22q11.2DS to behaviorally compensate for less efficient neural processing (Gothelf et al., 2007). In a cross-sectional study, response inhibition was shown to be impaired in older children with 22q11.2DS, but only in terms of accuracy (the outcome measure was proportion, more affected by bias than d-prime, a measure of sensitivity), not in the time taken to respond (Shapiro et al., 2013). Such inconsistencies may reflect methodological differences between studies and the significant phenotypic heterogeneity that is characteristic of 22q11.2DS (McDonald-McGinn et al., 2015; Philip & Bassett, 2011; Yamagishi et al., 1998). Additionally, while the two studies alluded to focused on children (Shapiro et al., 2013) and adolescents (Gothelf et al., 2007), here, we tested older individuals (adolescents and adults) with potentially more severe phenotypes. Interestingly, both of the current 22q11.2DS groups showed poorer d-prime than neurotypical controls, with their performance being more comparable to those of the schizophrenia group. That is, regardless of the presence of psychotic symptomatology, individuals with 22q11.2DS did not perform differently from individuals with schizophrenia, albeit the age differences in this comparison. This and previous evidence of poorer d-prime scores in unaffected siblings of individuals with schizophrenia (Groom et al., 2008) suggest that, in this context, d-prime might be indexing risk for psychosis, rather than exclusively disease-related processes. Both 22q11.2DS groups were faster than the schizophrenia group. However, whereas 22q11.2DS-did not differ from the neurotypical controls, individuals in the 22q11.2DS+ group were significantly slower than their neurotypical peers. This could suggest that, in 22q11.2DS, as in schizophrenia (Jogems-Kosterman, Zitman, Van Hoof, & Hulstijn, 2001), psychomotor slowing might be more pronounced in those with more severe phenotypes.

EEG analyses focused on components associated with different elements of response inhibition. In the context of Go/No-Go tasks, while the N2 has been argued to index early, automatic inhibition (De Sanctis et al., 2014; Eimer, 1993; Malcolm et al., 2015; O’Connell et al., 2009) and/or conflict detection processes (Dockree et al., 2005; Donkers & Van Boxtel, 2004; Morie et al., 2014), the P3 has been theorized as a marker of response inhibition (Bokura et al., 2001; Eimer, 1993; Enriquez-Geppert, Konrad, Pantev, & Huster, 2010; Groom & Cragg, 2015; Kiefer et al., 1998; Waller et al., 2019; Wessel & Aron, 2015), stimulus evaluation (Benvenuti et al., 2015; Bruin & Wijers, 2002; Smith et al., 2008) and adaptive, more effortful forms of control (De Sanctis et al., 2014; Malcolm et al., 2015; Wiersema & Roeyers, 2009). Our findings indicate that, in the N2 time window, and as previously shown, differences between hits and correct rejections are reduced in schizophrenia (Groom et al., 2008; Kiehl, Smith, et al., 2000) (but see (Weisbrod et al., 2000) for evidence of an intact N2 but impaired P3 in this population). However, no differences were found between 22q11.2DS and neurotypical controls, possibly reflecting the lack of a clear N2 effect in either group. The N2 has been argued as a less reliable marker of response inhibition than the P3 (Kropotov, Ponomarev, Hollup, & Mueller, 2011; Randall & Smith, 2011; Smith, Johnstone, & Barry, 2007) and, in fact, here, P3 was significantly reduced, not only in schizophrenia, but also in 22q11.2DS. Interestingly, Figure 6 seems to suggest that while a No-Go P3 was not reliably elicited in schizophrenia, though reduced, it was elicited in the 22q11.2DS group. Thus, it would appear that P3 is indexing severity of disease—while those with 22q11.2DS are at risk or at early stages of the illness, the individuals with schizophrenia in this study are chronic. And indeed, P3 reductions in schizophrenia (Groom et al., 2008) have been associated with disease severity, regardless of medication intake and task demands (Pfefferbaum, Ford, White, & Roth, 1989) (but see (Mathalon, Ford, & Pfefferbaum, 2000) for evidence that only auditory P3 is a trait marker for schizophrenia). However, P3 reductions have been likewise shown in other conditions such as, for instance, chronic alcoholism (Kamarajan et al., 2005) and ADHD (Fisher, Aharon-Peretz, & Pratt, 2011), suggesting that decreases in the No-Go P3 may index general cognitive impairment in conditions characterized by inhibition deficits, rather than schizophrenia-specific abnormalities. The No-Go P3, might, nevertheless, be a good indicator of disease severity and may have potential to differentiate, within the 22q11.2DS population, those at higher risk to develop schizophrenia. And, remarkably, when considering the 22q11.2DS groups separately, while in 22q11.2DS+ the P3 was reduced compared to neurotypical controls, but did not differ from those with schizophrenia; in 22q11.2DS-the P3 was increased when compared to those with schizophrenia, but did not differ from neurotypical controls. Furthermore, the correlations performed suggested associations between P3 and the D-KEFS inhibition measure: Those with smaller P3s performed worse in the D-KEFS inhibition task, which further argues for the No-Go P3 as reflecting the ability to inhibit a prepotent response. P3 appears to be associated with activations in the inferior frontal cortex (IFC) (Enriquez-Geppert et al., 2010). Impaired white matter integrity in the IFC is reduced in schizophrenia (Buchanan, Vladar, Barta, & Pearlson, 1998) and has been associated with severity of negative symptoms (Wolkin et al., 2003). In 22q11.2DS, the IFC seems to be characterized by increased cortical thickness (Jalbrzikowski et al., 2013). Differences in cortical thickness in this population have been associated with cognitive abilities in children and adolescents and schizophrenia in adults (Schaer et al., 2009). P3 has additionally been argued to reflect activity of the neuromodulatory locus coeruleus (LC)–norepinephrine (NE) producing nucleus (Nieuwenhuis, Aston-Jones, & Cohen, 2005). Interestingly, the IFC might be particularly influenced by NE release if LC neurons are activated by the decision to inhibit a response (Aston-Jones & Gold, 2009). Though anatomically the LC—which comprises the largest cluster of norepinephric neurons in the brain—does not seem to, at least as measured post-mortem, differ between individuals with schizophrenia and controls (Craven, Priddle, Crow, & Esiri, 2005), norepinephric dysfunction has been shown in schizophrenia (Yamamoto & Hornykiewicz, 2004) and associated with the cognitive deficits characteristic of the disorder (Friedman, Adler, & Davis, 1999). COMT, which encodes the protein catechol-O-methyltransferase responsible for degrading catecholamines such as norepinephrine (particularly in the prefrontal cortex), is a gene in the 22q11.2 region, and is often alluded to as a plausible candidate for psychiatric disorders (Philip & Bassett, 2011).

As observed for the P3, 22q11.2DS and schizophrenia groups presented reduced Ne and Pe amplitudes. No clear consensus has been reached on the functional interpretation of the Ne, though its attenuation has been associated with reduced activation of the rostral and caudal anterior cingulate (Laurens, Ngan, Bates, Kiehl, & Liddle, 2003; Veen & Carter, 2002). While some evidence indicates that Ne amplitudes are larger when one is aware that an error has been made (Luu, Flaisch, & Tucker, 2000), others suggest that the Ne is not modulated by the awareness that an error was committed (Hester et al., 2005; Nieuwenhuis et al., 2001). Nonetheless, Ne is consistently reported as attenuated in individuals with schizophrenia (Alain, McNeely, He, Christensen, & West, 2002; A. T. Bates, Kiehl, Laurens, & Liddle, 2002; Houthoofd et al., 2013; Kopp & Rist, 1999; Mathalon et al., 2002) and has been argued as a potentially important trait of the illness, evident well before the sequelae emerge. Decreased Ne amplitudes have been shown in children with putative antecedents of schizophrenia (Laurens et al., 2010) and in chronic, early illness and high-risk individuals (Perez et al., 2012). Consistent with such findings, here, Ne was reduced not only in the schizophrenia group, but also in 22q11.2DS, an at-risk population. While those with 22q11.2DS but no psychotic symptoms did not differ, however, from the neurotypical controls, we observed no differences between 22q11.2DS- and 22q11.2DS+ and either of the 22q11.2DS groups and schizophrenia. Larger samples of individuals with 22q11.2DS with and without psychotic symptoms would be needed to better understand the absence of differences between 22q11.2DS- and neurotypical controls. Nevertheless, our findings are consistent with the suggestion of Ne as a marker of risk for psychosis.

Regarding the Pe, while some have shown reductions of this component in schizophrenia (Foti, Kotov, Bromet, & Hajcak, 2012; Perez et al., 2012), others have not found differences between individuals with schizophrenia and neurotypical controls (Alain et al., 2002; Kim et al., 2006; Mathalon et al., 2002; Morris, Yee, & Nuechterlein, 2006). It is worth noting that some of the studies that found no differences (Kim et al., 2006; Morris et al., 2006) used relatively conservative high-pass filters (1-2 Hz), which could have filtered out Pe entirely. Pe reductions suggest that these clinical populations have a weakened (or even absent) sense of error awareness (O’Connell et al., 2007). The post-error slowing in 22q11.2DS and schizophrenia (reflected in longer reaction times in the trial after false alarm, Figure 2) in our data suggest, nevertheless, that those errors might have been, at least partially, acknowledged. Pe could thus additionally index a subjective or emotional error evaluation process, possibly modulated by the individual significance of the error (Falkenstein, Hoormann, Christ, & Hohnsbein, 2000). Individuals who commit errors more often (i.e., those with higher rates of false alarms), might attribute lower subjective or emotional significance to the errors made than those who rarely commit them. A lower attributed significance could thus result in a smaller Pe amplitude. Though this interpretation of Pe would further fit fMRI evidence suggesting that the rostral anterior cingulate, which is associated with affective processes, may be a key player during post-error activity (Kiehl, Liddle, & Hopfinger, 2000; Magno, Foxe, Molholm, Robertson, & Garavan, 2006), here, individuals with 22q11.2DS and those diagnosed with schizophrenia did not differ from the neurotypical controls in false alarms rates. And, as can be appreciated in Figure 2, neurotypical controls, despite their apparent awareness of the errors committed reflected on the increased Pe amplitude, did not present such a reaction time lengthening in the trial after false alarms, which might indicate that the post-error slowing observed in 22q11.2DS and schizophrenia is mostly reflective of slower processing of error information. Contrary to what was described for the P3 and similarly to what was seen in the Ne, no differences were found between 22q11.2DS- and 22q11.2DS+ in the Pe: Both groups showed similar amplitudes to the ones observed in the schizophrenia group, and clearly reduced when compared to the neurotypical group. Pe could, therefore, represent a marker of risk for schizophrenia.

Ne and Pe amplitudes were correlated with number of commission errors, where erroneous responses were made to the repeated image (non-targets), and with the D-KEFS inhibition measure, which confirms these components’ functional association with inhibition. The associations between Ne and Pe and the inhibition score were, nevertheless, exclusively dictated by the clinical groups, with no correlation observed in the neurotypical controls. As can be appreciated in Figure 13, this is likely explained by the greater variability observed in the clinical groups in the inhibition score, which suggests the potential usefulness of this measure to characterize populations with response inhibition difficulties. Moreover, these associations offer additional support to a recent study suggesting that performance monitoring impairment (as assessed through Ne and Pe) contributes to executive function deficits, which in turn contributes to negative symptoms in psychosis (Foti et al., 2020).

Lastly, though the focus of the current study was primarily on response inhibition, we expected alterations in early sensory processing in the clinical groups, given the reported differences in early visual-evoked potentials in schizophrenia (Butler & Javitt, 2005; Foxe et al., 2001; Foxe et al., 2005; Lalor, Yeap, Reilly, Pearlmutter, & Foxe, 2008; Yeap, Kelly, Sehatpour, et al., 2008; Yeap et al., 2006; Yeap, Kelly, Thakore, & Foxe, 2008) and in 22q11.2DS (Biria et al., 2018; Magnee et al., 2011). Basic visual sensory responses appeared slightly reduced in both clinical populations when compared to neurotypical controls. In 22q11.2DS, those differences were not, however, statistically significant. The few studies investigating visual processing in this population revealed patterns of increased and decreased amplitudes, depending on the time window of interest or on the task at hand (Biria et al., 2018; Magnee et al., 2011). The mixed nature of the current 22q11.2DS sample (individuals with and without symptoms) could have impacted the ability to detect differences between those with 22q11.2DS and their age-matched neurotypical peers. A recent EEG study focused on basic auditory processing in this population suggested differences between those with and without psychotic symptomatology and argued for the presence of two opposite mechanisms in those with the deletion: one that relates to a process specific to the deletion itself and gives rise to larger amplitudes, and another, associated with psychosis that results, as in schizophrenia, in decreases in amplitude (Francisco, Foxe, Horsthuis, DeMaio, & Molholm, 2020). And, indeed, here, whereas the 22q11.2DS- group did not differ from the neurotypical controls, but presented increased amplitudes when compared to the schizophrenia group; 22q11.2DS+ did not differ from the individuals with schizophrenia, but presented decreased amplitudes when compared to the neurotypical controls.

No differences were likewise found between individuals with schizophrenia and their neurotypical peers in our primary analysis. However, and in accordance with previous studies (Foxe et al., 2001; Yeap, Kelly, Sehatpour, et al., 2008), post-hoc statistical cluster plots suggested the presence of differences between the two groups in the P1 time window. Such differences might not have been captured by the more conservative model used in the primary analysis. The absence of statistically significant differences between the schizophrenia group and its age-matched controls could additionally be explained by the inclusion of two individuals with schizophrenia whose amplitudes in the time windows of interest were significantly larger than the remainder individuals part of that group. When those individuals—two of the youngest in the group, with somewhat recent diagnoses and thus a relatively short history of major symptomatology and medication use—were excluded from the group, statistically significant differences were found between the individuals with schizophrenia and their age-matched peers in the P1 and P2. A larger sample including individuals at different stages of the illness would be needed to better characterize the impact of illness chronicity and medication use in basic visual processing. The early visual processing components assessed here seem to reflect disease, not risk, since no difference were found between those with 22q11.2DS but no psychotic symptoms (though still at genetic risk) and the neurotypical control group. This would appear to contrast with Yeap et al (2006), in which reduced visual P1 was demonstrated in unaffected first-degree relatives of individuals with schizophrenia as well as probands. One possibility is that the data from the 22q11.2DS sample that we tested here may reflect opposing effects on sensory processing, which combine to mask P1 deficits associated with risk for psychosis. The correlations described between the visual components and the response inhibition markers suggest that differences in the extraction of information from early visual processing may have implications for information processing at later stages.

Some limitations to this study should be noted. Despite the satisfactory size of our sample considering the nature of any rare disease, larger numbers would permit more detailed analyses, such as those focused on associations between neural, cognitive, and behavioral outcomes. Furthermore, they would allow one to take into consideration the impact of medication and of co-morbidities. Additionally, though findings point toward a striking similitude between individuals with 22q11.2DS and psychotic symptoms and individuals with schizophrenia regardless of the age differences between these two groups, ideally the groups should be age matched. A younger schizophrenia group would also be important to reduce confounds associated with long-term antipsychotic use, recurrent psychotic episodes, and institutionalization. In future work, additional measures of psychotic symptoms should be used to more fully characterize psychosis in these populations: Measures providing symptom severity and the differentiation between negative and positive symptoms could be more informative.

In summary, this study provides the first EEG evidence of inhibitory deficits in 22q11.2DS and pinpoints the stages of information processing that are affected. It additionally adds to the extant literature by showing differences between those with and without psychotic symptoms, which contributes to a better characterization of this syndrome and of those at-risk for psychosis not only in 22q11.2DS, but also, potentially, in the general population. We have additionally showed that whereas P3 may be useful as marker of disease severity, Ne and Pe might be biomarkers of risk for schizophrenia. That the differences between those with 22q11.2DS with and without psychotic symptoms were much more noticeable and consistent when using electrophysiology (when compared to the behavioral measures), indicates the potential of electrophysiological methods as powerful approaches to early detection. Given how crucial inhibitory processes are to cognitive and daily function, our findings have the potential to inform interventions to improve cognitive control and overall functioning and the development of self-management strategies in both 22q11.2DS and schizophrenia.

## ACKNOWLEDGEMENTS

We wish to thank Dr. Juliana Bates, who performed the clinical assessments, and Elise Taverna and Danielle Newbury for their help with data collection. We would also like to acknowledge the role of the Jacobi Medical Center and of the Montefiore-Einstein Regional Center for 22q11.2 Deletion Syndrome in recruitment, and the Rose F. Kennedy Intellectual and Developmental Disability Research Center for all of its support. We extend our most sincere gratitude to the participants and their families for their interest, their involvement, and their time. This work was supported in part by the Eunice Kennedy Shriver National Institute of Child Health and Human Development (NICHD), under award number U54 HD090260.

## AUTHOR CONTRIBUTIONS

SM, AAF, and JJF conceived the study. AAF and DJH collected and analyzed the data. AAF wrote the first draft of the manuscript. SM, JJF, and DJH provided editorial input to AAF on the subsequent drafts.

## SUPPLEMENTARY FIGURES

**Supplementary Figure 1.**
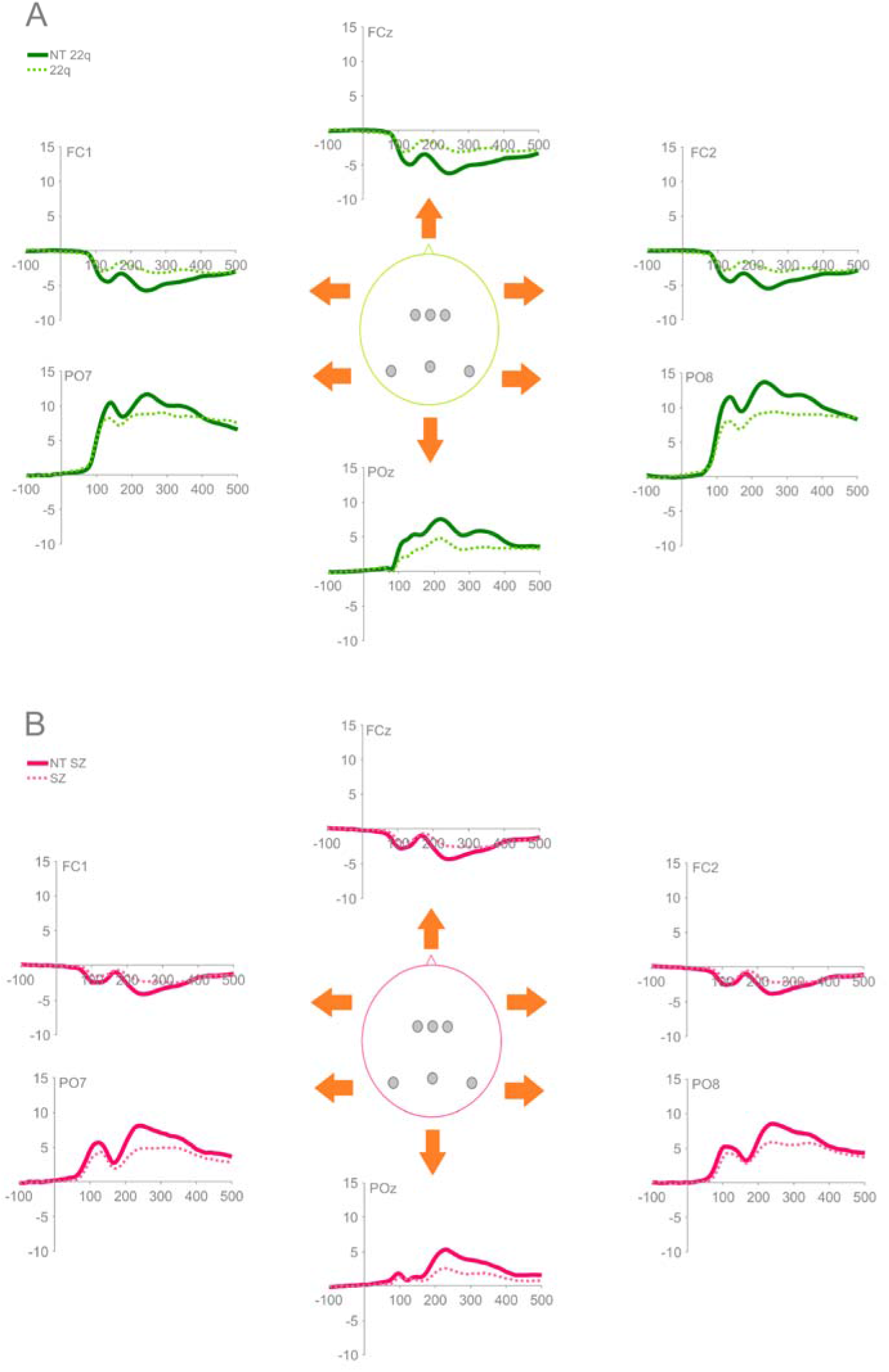
Averaged ERPs for the early sensory time windows of interest (P1, N1, and P2), from channels over left (FC1), center (FCz), and right (FC2) frontocentral and left (PO7), center (POz), and right (PO8) posterior scalp regions for NT-22q (Panel A) and NT-SZ (Panel B).

**Supplementary Figure 2.**
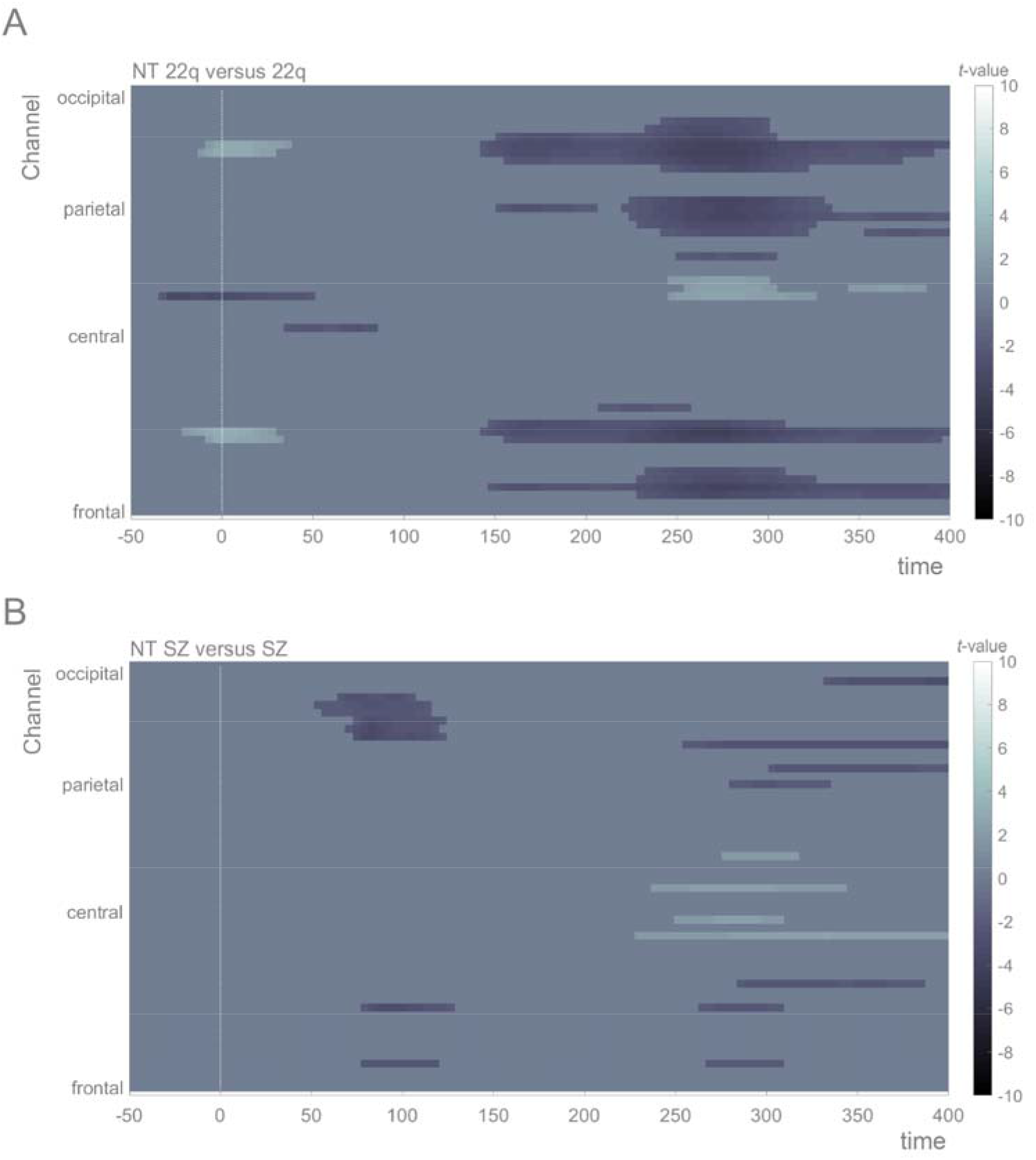
Statistical cluster plots of the sensory responses. Color values indicate the p-values that result from point-wise *t*-tests evaluating the visual responses across time (*x*-axis) and electrode positions (*y*-axis). General electrode positions are arranged from frontal to occipital regions (bottom to top) and the scalp has been divided into four general scalp regions. Within each general region, electrode laterality is arranged from left to right Only *p*-values <0.05 are color-coded.

